# Seeing Through Touch: Visual Reliability Shapes Tactile Guidance in 360° Virtual Reality

**DOI:** 10.64898/2026.08.10.744038

**Authors:** Ailene Y. C. Chan, Shinsuke Shimojo

## Abstract

This study characterizes how people combine visual and tactile directional cues while acting in a fully immersive 360° virtual environment. Participants used a vibrotactile belt and VR headset to localize targets while we manipulated visual reliability and the spatial discrepancy between visual and tactile signals. Behaviorally, degraded visual input made visual responses slower, less precise, and more susceptible to tactile pull, whereas tactile-guided responses remained comparatively stable.

We then asked whether these behavioral changes reflected a change in multisensory binding or a change in sensory uncertainty. A Bayesian Causal Inference (BCI) framework captured the structure of behavior under high visual reliability and continued to track individual differences under low visual reliability, even though its absolute goodness-of-fit decreased. Under extreme visual noise, Bayesian Information Criterion sometimes favored a simpler Maximum Likelihood Estimation (MLE) model, but MLE showed poor absolute fit and did not capture meaningful behavioral variability. This dissociation shows that statistical parsimony and explanatory validity can diverge when behavior becomes highly variable.

BCI-derived parameters further indicated that degraded vision increased visual uncertainty, while the prior tendency to bind visual and tactile cues remained stable. Kinematic analyses added a complementary insight: early movement trajectories were strongly shaped by tactile signals, even when final localization was visually guided. Together, these findings suggest that visual–tactile integration in 360° environments depends on sensory reliability and task demands, with tactile cues providing fast body-centered guidance when visual information is limited.

## 1 Introduction

Navigating complex 360-degree environments requires the nervous system to combine sensory signals that are often noisy, incomplete, or spatially discrepant. Virtual reality (VR) provides a controlled way to study this problem while preserving the demands of active spatial behavior (Marucci et al., 2021). In head-mounted displays, however, objects outside the current field of view often require extensive visual search through head rotation, which can slow performance and increase discomfort (Ariza N. et al., 2017). Vibrotactile displays worn on the torso offer a complementary channel for omnidirectional cueing because the torso provides a natural egocentric map of external direction (Terrence et al., 2005; Choi and Kuchenbecker, 2013; Cholewiak et al., 2004; van Erp, 2008). When visual and tactile cues are both available, the nervous system must decide whether they should be bound into a common percept or treated as independent events.

Bayesian Causal Inference (BCI) provides a formal framework for this decision. In BCI models, observers estimate whether cues share a common cause and combine them according to their relative reliability (Ernst and Banks, 2002; Körding et al., 2007; Shams and Beierholm, 2022; Wozny et al., 2010). This framework has been studied extensively in visual– auditory spatial tasks, but those experiments have largely used two-dimensional displays or restricted frontal fields of view (Beierholm et al., 2009; Rohe and Noppeney, 2015; Aller and Noppeney, 2019; Odegaard et al., 2017; Odegaard and Shams, 2016; Odegaard et al., 2016). Computational studies of visual–tactile integration are less common and have similarly focused on frontal space (Verhaar et al., 2022) or non-spatial domains such as numerosity (Wozny et al., 2008). It is still unclear what happens when vision becomes unreliable during an active VR task, especially when visual and tactile cues can point to different locations around the body.

This question matters for both basic theory and applied systems. For people with low vision or blindness, understanding how degraded visual input is weighted against tactile information can inform sensory substitution and navigation devices (Paterson, 2021; Eklund and Helmefalk, 2018; Spence, 2014). Similar principles apply to sighted users operating in visually uncertain environments, where spatial tactile displays may provide directional information without further burdening visual attention (Marucci et al., 2021; Elliott and Redden, 2012). Yet applied studies of vibrotactile cueing often focus on performance outcomes rather than the computational mechanisms that determine when tactile signals are integrated, ignored, or used to guide action (de Jesus Oliveira et al., 2017; Van Erp, 2005; Rahal et al., 2009; Luzhnica et al., 2017).

The present study investigates visual–tactile integration in a 360-degree VR targeting task using a custom vibrotactile belt with eight eccentric-rotating-mass tactors. We manipulated two factors: the spatial discrepancy between visual and tactile targets, and the reliability of visual input. Behavioral analyses tested performance across the full 360-degree space, whereas computational modeling focused on front-region trials where the visual reliability manipulation most directly constrained sensory inference. This design allowed us to ask three linked questions. First, does degraded vision increase tactile capture of visual responses? Second, do movement trajectories reveal action strategies that differ from the final perceptual endpoint? Third, does visual uncertainty change the tendency to bind visual and tactile cues, or does it primarily alter sensory noise and priors? By combining accuracy, directional bias, kinematic classification, and BCI model fitting, we aim to separate changes in sensory uncertainty from changes in the underlying rules of multisensory binding.

## 2 Methods

### 2.1 Human participants

Twenty-five participants (15 females; range = 18-53 years, mean = 30.1, SD = 7.5) took part in the study. All participants had normal or corrected-to-normal (20/20) vision. With the exception of one author (*N* = 1), all participants were naïve to the purpose of the study. Written informed consent was obtained from all participants prior to the session. Participants were compensated at a rate of $25/hour. All procedures were approved by the Institutional Review Board at the California Institute of Technology (Protocol IR24-0926).

### 2.2 Materials and apparatus

#### 2.2.1 Vibrotactile Belt

Directional tactile cues were delivered via a custom-built vibrotactile belt constructed from a loop-style inner duty belt (Fig. 1A). The system featured eight miniature eccentric-rotating-mass (ERM) tactors (Pico Vibe 9 mm, 25 mm type; model 307–103; Precision Microdrives, London, UK) controlled by an Arduino Nano 33 BLE Sense Rev2. The tactors were driven by a custom printed circuit board (PCB) designed by D. Wagenaar and A. Chan and assembled in-house (Fig. 1B).

**Figure 1:**
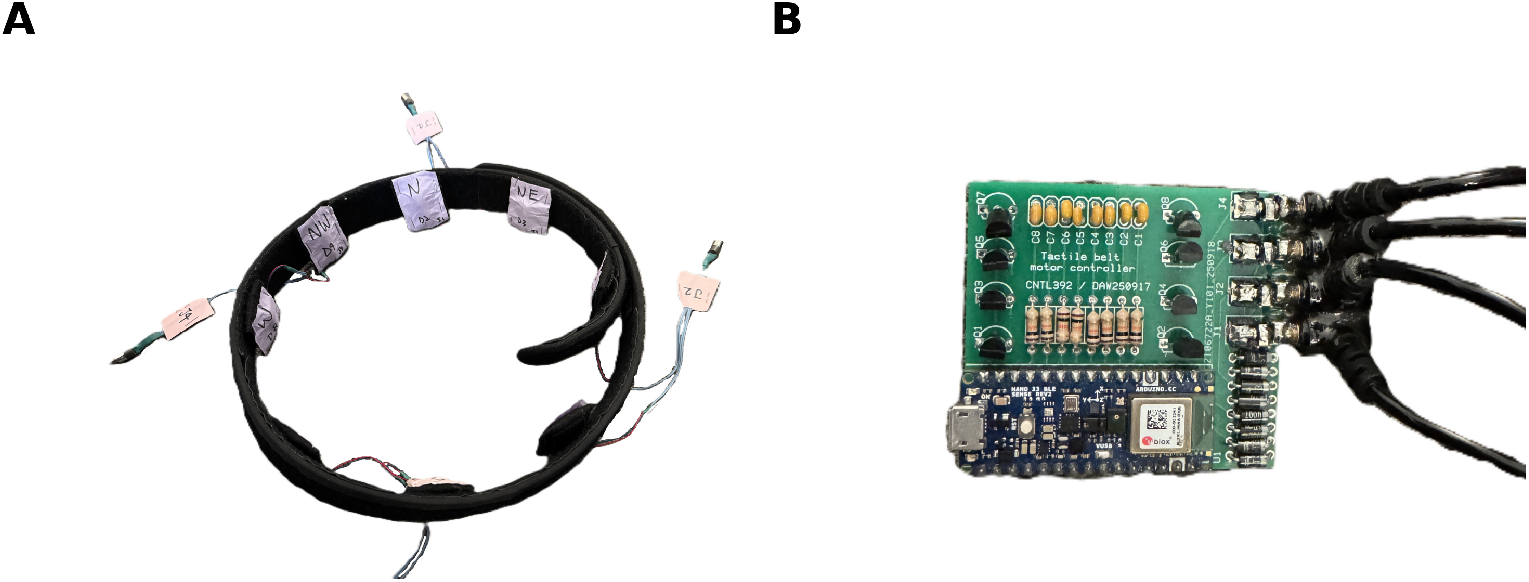
Vibrotactile Belt Prototype and Control Hardware. (A) The wearable belt assembly featuring eight tactors distributed at cardinal and collateral positions. (B) The custom PCB interface integrated with an Arduino Nano 33 BLE Sense Rev2 for signal processing and wireless communication.

Each tactor was housed in an individual felt pocket with Velcro backing, allowing the tactors to be repositioned easily to accommodate different waist sizes. To prevent rotation of tactors inside the felt pocket, which would compromise wire integrity, D. Wagenaar designed and 3D-printed tactor holders to restrict tactor rotations. Tactors were aligned with the participant’s cardinal (N, E, S, W) and collateral (NE, NW, SE, SW) points relative to the torso. Detailed circuit schematics and PCB layouts are provided in Fig. A.1.

#### 2.2.2 Virtual Reality Headset

Visual stimuli were presented using a Meta Quest 2 head-mounted display (HMD). Participants interacted with the environment using a single right-hand VR controller to aim and “shoot” at targets.

#### 2.2.3 Software

The experimental task was developed in Unity (Unity Technologies) using custom C# scripts. The virtual environment placed participants at the center of an open space (Fig. 2A) where they scanned the horizon for Unidentified Flying Objects (UFOs; Fig. 2B) to engage with a virtual laser gun (Fig. 2C). 3D assets were sourced from Sketchfab under a standard license. Representative gameplay and trial procedures are illustrated in Supplementary Video 1.

**Figure 2:**
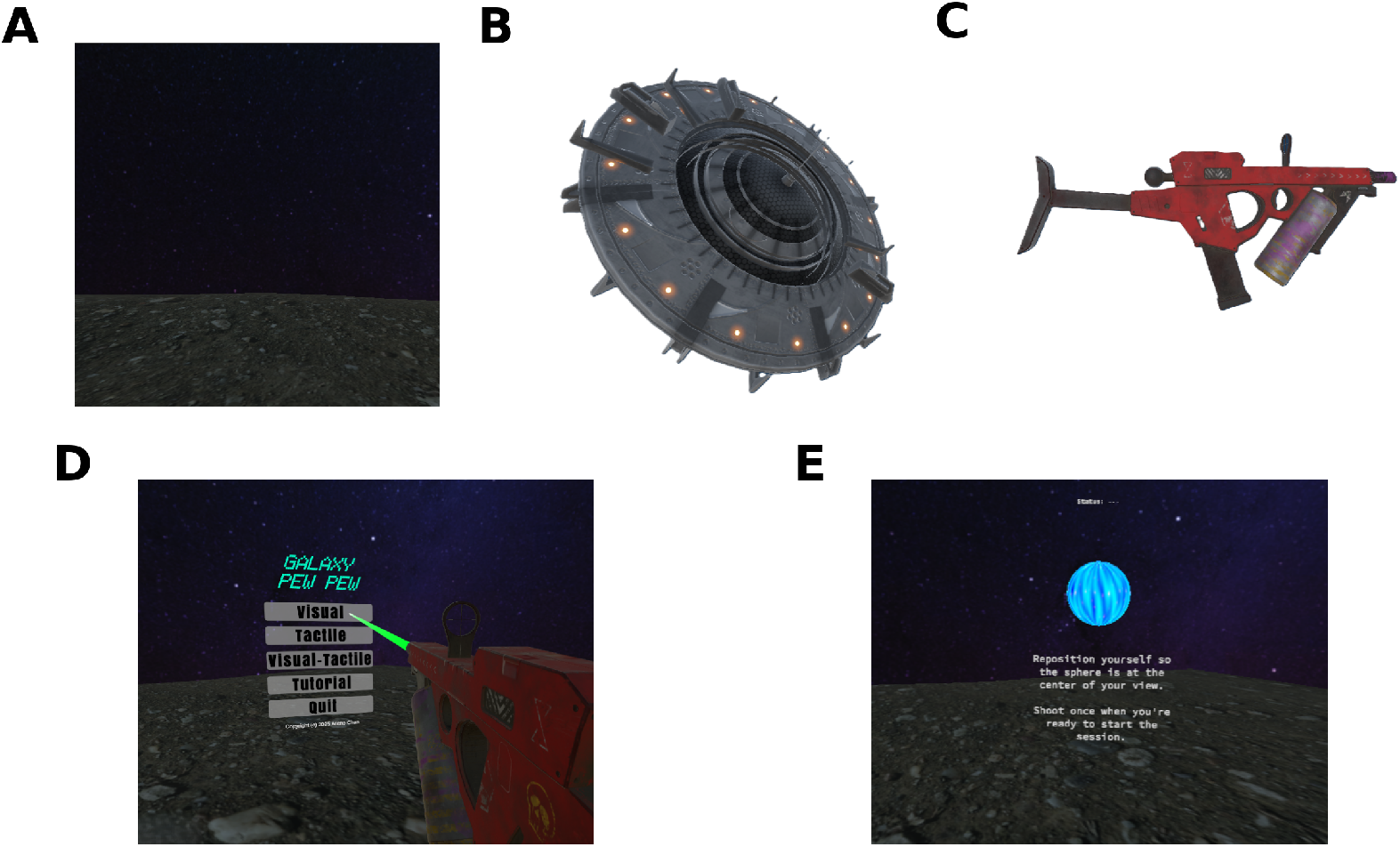
Visual assets and integrated environment. (A) Base terrain showing surface texture and starfield background. (B) Unidentified Flying Objects (UFO) model. (C) Laser gun used for player interaction. (D) Final composite scene of the main menu interface demonstrating the integration of all assets within the game engine. (E) Calibration scene.

Communication between the Unity application and the Arduino-based belt was established via Bluetooth Low Energy (BLE) using the *Bluetooth LE for iOS, tvOS and Android plugin* (Shatalmic, LLC). The software managed trial randomization, target positioning, and tactor activation. It recorded behavioral data at 60 Hz, including shooting accuracy (angle shot, angular error), reaction time, and continuous heading (yaw) for both the HMD and the handheld controller.

### 2.3 Design and procedure

The experimental task was a spatial shooting paradigm implemented in a fully immersive virtual environment. Participants were positioned at the center of an open 360° space and were instructed to remain seated, but they were free to move their upper bodies to scan the environment and execute one shot on each trial.

The task included three block types. In **unimodal visual-only (V) blocks**, one visual target appeared at a random direction at eye level, no tactile stimulus was presented, and participants shot at the visual target. In **unimodal tactile-only (T) blocks**, no visual target was presented; instead, one tactor vibrated, and participants shot in the direction indicated by the vibration. In **bimodal visual–tactile (VT) blocks**, both a visual target and a tactile stimulus were presented. On visual-response trials, participants shot at the visual target; on tactile-response trials, participants shot at the tactile target. The visual and tactile stimuli could either match or conflict, allowing us to measure how much one modality biased responses to the other.

On each trial, the stimulus or stimulus pair was presented for 3 seconds, after which participants were required to respond regardless of whether the target had been successfully localized.

Auditory cues were included for environmental ambiance and trial timing. Specifically, an alarm sound signaled the onset of each trial when the target appeared. However, all sounds were non-spatialized (2D) and did not convey directional information.

The session consisted of six blocks: two unimodal baseline blocks followed by four bimodal blocks. Participants always completed the unimodal Visual (V) and Tactile (T) blocks first to establish baseline performance. At the start of each block, participants selected the corresponding block type (V, T, or VT) or a tutorial from the Main Menu Scene (Fig. 2D). This was followed immediately by a Calibration Scene (Fig. 2E), allowing participants to physically reposition themselves so their “true North” aligned with the in-game orientation. Once calibrated, participants entered an Instruction Scene and initiated the experiment when ready. A total of eight directions were tested: cardinal (N, E, S, W) and collateral (NE, NW, SE, SW). For analysis, these eight directions were further categorized into three regions: front (N, NE, NW), side (E, W), and back (S, SE, SW) (Fig. 3A).

**Figure 3:**
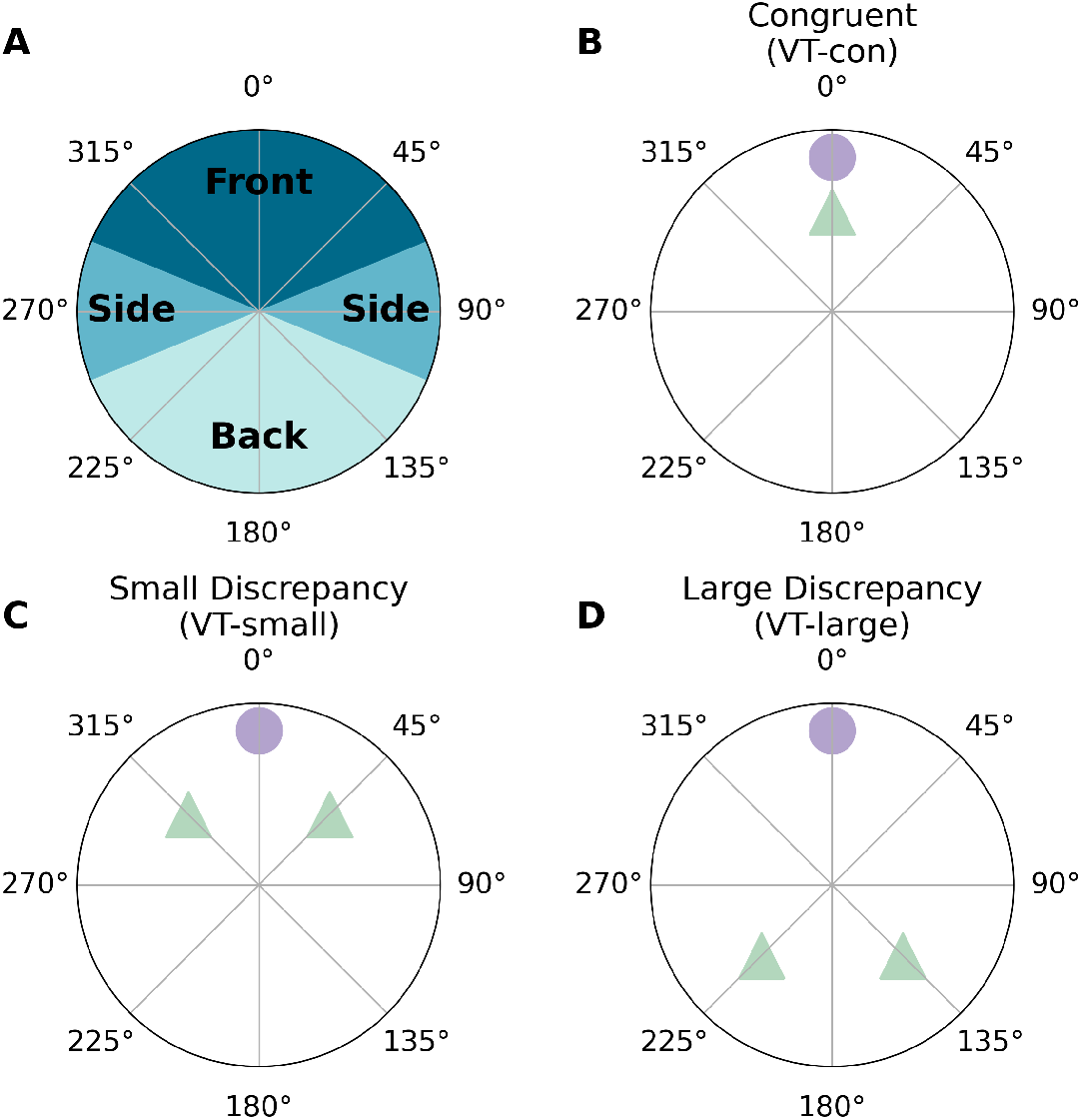
Visual and Tactile Stimulus Directions. (A) Stimulus directions categorized into three regions: front, side, and back. (B–D) Visual targets (purple circles) and tactile cues (green triangles) in VT-congruent, VT-small, and VT-large conditions.

The subsequent multisensory Visual-Tactile (VT) blocks were grouped by required response modality (Visual or Tactile). To mitigate fatigue, each response modality condition was split into two blocks. Within VT blocks, trials were categorized into three spatial subconditions: (1) **Congruent** (VT-congruent): 0° spatial offset between visual and tactile stimuli (Fig. 3B), (2) **Small Discrepancy** (VT-small): *±*45° offset (e.g., a 0° visual target paired with a NW or NE tactor; Fig. 3C), and (3) **Large Discrepancy** (VT-large): *±*135° offset (e.g., a 0° visual target paired with a SW or SE tactor; Fig. 3D). For both discrepancy conditions, the direction of the offset (positive or negative) was randomized across trials within each block.

Visual reliability manipulation was achieved by overlaying a spherical mask over the main camera (participants’ first person point of view) and adjusting its alpha (opacity) in Unity: **High Reliability** (alpha = 0) and **Low Reliability** (alpha = 245; max 255) (Fig. 4). The alpha value for the low reliability condition was selected to maximize the contrast between conditions within the technical constraints of the headset. Although higher opacity values were tested, alpha = 245 represented the highest level at which the visual target remained faintly visible, avoiding complete disappearance due to display resolution limits.

**Figure 4:**
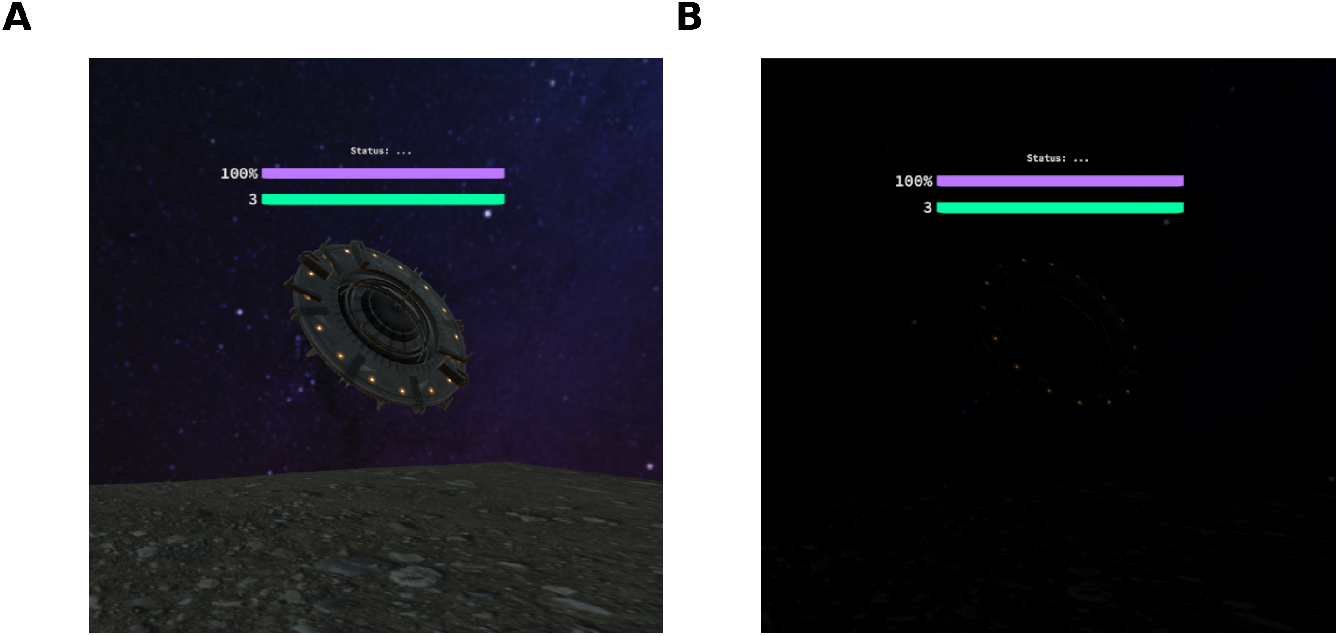
Visual Reliability Levels. (A) High (alpha = 0), (B) Low (alpha = 245).

The unimodal V block followed a 2 *×* 8 *×* 5 factorial design (Visual Reliability *×* Angle *×* Repetition), totaling 80 trials. The unimodal T block followed an 8 *×* 5 factorial design (Angle *×* Repetition) for 40 trials. Finally, the bimodal VT blocks followed a 2 *×* 8 *×* 3 *×* 5 factorial design incorporating visual reliability, angle, spatial subcondition, and repetitions.

### 2.4 Data analysis

#### 2.4.1 Accuracy

Accuracy was defined as the spatial proximity of the participant’s shot angle to the visual or tactile target. To account for the 360° spherical space, we first calculated the circular difference, Δ*θ*, between the shot angle (*θ*_shot_) and the target angle (*θ*_target_), constrained to the range [*−*180, 180]°. Accuracy was then normalized to a [0, 1] scale using the following transformation:

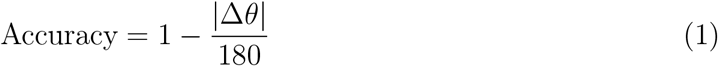

In this metric, an accuracy of 1.0 represents a perfect hit (zero angular error), while an accuracy of 0.5 represents a shot fired at a 90° offset. This normalized approach facilitates a direct comparison across different conditions and target directions.

Data were analyzed using Linear Mixed Models (LMM) to account for the nested structure of trials within participants. For all models, Participant ID was included as a random intercept to control for individual differences in baseline spatial performance.

To evaluate multisensory performance, we first conducted a 3-way LMM with Condition (Spatial Disparity: Congruent, Small, Large), Response Type (Visual, Tactile), and Visual Reliability (High, Low) as fixed effects. To investigate the potential impact of spatial region (Front, Side, Back) on performance, a second 4-way LMM was performed, incorporating Region as an additional fixed factor.

#### 2.4.2 Directional Bias and Sensory Conflict

To quantify the degree to which the tactile distractor pulled the participant’s response away from the visual target, we calculated the mean directional bias 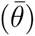 using circular statistics. For each condition, the angular errors were wrapped to the range [*−*180, 180]° and the circular mean was computed:

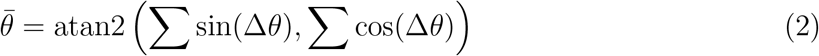

A bias of 0° indicates a response centered on the target, while a bias shifting toward the distractor angle (45° or 135°) indicates multisensory capture. Additionally, we calculated the vector strength (*R*), also known as the resultant length, to measure response precision:

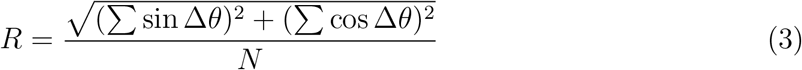

The *R* value ranges from 0 to 1, where 1.0 represents perfect directional consensus among trials and 0 represents a uniform random distribution. Significant differences in mean directional bias between reliability conditions were assessed using the Watson-Williams test, a circular analogue of a one-way ANOVA.

#### 2.4.3 Movement Trajectories

##### Visualization

To compare trajectories across conditions and subjects with varying reaction times (RT), movement data were standardized using a retrospective temporal alignment. For each trial, raw yaw values for the head and controller were unwrapped to a continuous angular scale and smoothed using a centered moving average filter (window = 20 frames; 60 Hz). Movement error was defined as the angular distance between the effector and the target, wrapped within [*−*180*^◦^,* 180°]. To account for varying trial durations, trajectories were clipped from the start of trial (*t_start_*) to the moment of the “shot” (*t_shot_*). All trials were then interpolated onto a common 5-second temporal grid relative to the end of the trial (*t_shot_*), allowing for subject-wise and group-level averaging.

##### Random Forest Classification

We used Random Forest classifiers to test whether movement patterns could identify what kind of trial participants were doing. The classifiers used movement features only; final angular error was excluded so that the analysis captured *how* participants moved, rather than whether the final shot was accurate. For each trial, we extracted four features from both the head and controller trajectories (8 features total):

- **Max Velocity (***v_max_***, deg/s):** how fast the movement became.
- **Peak Acceleration (***a_peak_***, deg/s**^2^): how sharply the movement started or changed speed.
- **Mean Jerk (***j̅***, deg/s**^3^): how smooth or abrupt the movement was.
- **Path Length (***L***, deg):** how much the head or controller moved before the shot.

All derivatives were computed using the unwrapped angular trajectories to avoid artifacts from circular wrap-around points.

We trained separate classifiers for five questions:

1. **Unimodal Baseline (V vs. T):** Can movement distinguish visual-only from tactile-only trials?
2. **Task Intent (Visual vs. Tactile response):** In multisensory trials, can movement reveal which cue participants were asked to report?
3. **Multisensory Conflict (Congruent vs. Small vs. Large):** Can movement reveal the size of the conflict between visual and tactile cues?
4. **Multisensory Enhancement (V vs. T vs. VT-congruent):** Do congruent visual– tactile trials look more like visual-only or tactile-only trials?
5. **Visual Reliability (High vs. Low):** Can movement reveal whether the visual target was easy or hard to see?

##### Model Architecture and Evaluation

Each classifier used 100 trees with a maximum depth of 10. Data were split into 70% training and 30% test sets using stratified sampling, and balanced class weights were used when trial counts differed across classes. A fixed random seed (42) was used for reproducibility. We report classification accuracy and F1 score, and used Gini importance to identify which movement features contributed most to each classifier.

#### 2.4.4 Model Fitting

We modeled multisensory integration using a Bayesian Causal Inference (BCI) framework to estimate the probability that visual (*V*) and tactile (*T*) cues originated from a common source (*C* = 1) or independent sources (*C* = 2). This framework allows us to derive forced-fusion, full-segregation, and maximum likelihood estimation (MLE) models as nested alternative hypotheses (Ernst and Banks, 2002; Körding et al., 2007; Wozny et al., 2010).

The BCI model assumes that visual and tactile stimuli at true spatial locations *s_V_*and *s_T_* generate noisy internal observations *x_V_* and *x_T_*. These sensory observations are modeled as independent samples drawn from Gaussian distributions centered on the true stimulus positions, with sensory noise defined by standard deviations *σ_V_* and *σ_T_* :

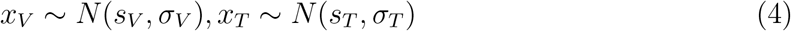

Furthermore, the model incorporates a spatial prior representing the observer’s expectations about stimulus locations. This prior is modeled as a Gaussian distribution centered at *µ_P_* with standard deviation *σ_P_* :

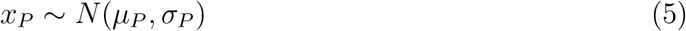

The observer must infer the underlying causal structure (*C*) that is hidden from the nervous system. The posterior probability of a common cause (*C* = 1) is calculated by combining the sensory evidence with a causal prior, *p_common_*, according to Bayes’ rule:

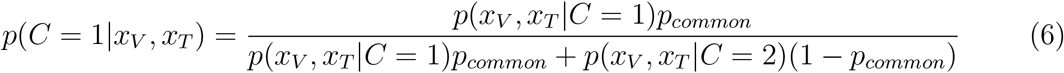

Here, *p_common_* represents the observer’s a priori belief that the visual and tactile signals belong to the same origin. Post-stimulus, the likelihood terms *p*(*x_V_, x_T_ |C*) quantify how likely the noisy internal observations *x_V_* and *x_T_* are, given each causal hypothesis.

If the signals are attributed to a common cause (*C* = 1), the optimal estimate of the stimulus location, spatial prior: *ŝ*_*C*=1_, is a reliability-weighted average of both sensory signals and the

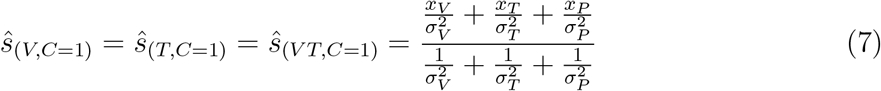

If the signals are attributed to independent causes (*C* = 2), the estimates for each modality are computed separately. In this case, each estimate is influenced only by its corresponding sensory signal and the spatial prior:

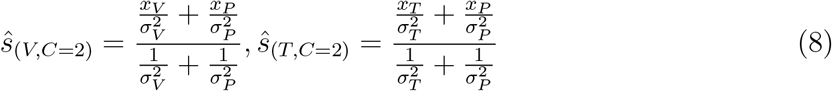

To obtain a final estimate of visual or tactile target location, we assumed the observer combines the estimates from the two causal structures (*ŝ*_*C*=1_ and *ŝ*_*C*=2_) using one of three common decision strategies (Wozny et al., 2010):

##### Model Averaging (MA)

The final estimate is a weighted sum of the estimates from both causal structures, where the weights are determined by their respective posterior probabilities:

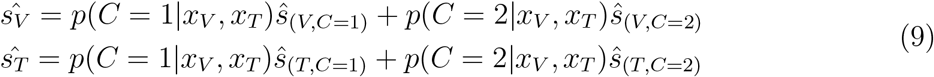

##### Model Selection (MS)

The observer selects the estimate from the causal structure with the higher posterior probability. If *p*(*C* = 1*|x_V_, x_T_*) *>* 0.5, the observer adopts *ŝ*_*C*=1_; otherwise, they adopt *ŝ*_*C*=2_:

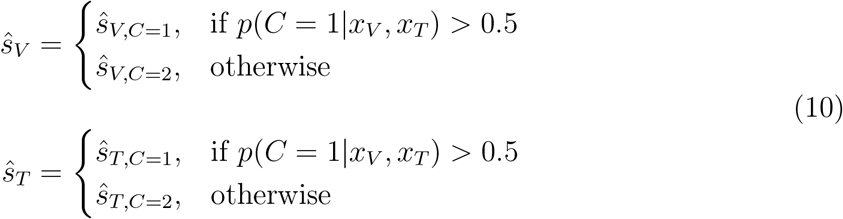

##### Probability Matching (PM)

A stochastic strategy where the observer chooses a structure proportional to its posterior probability. On each trial, a threshold *ξ* is drawn from a uniform distribution *U* (0, 1). If *p*(*C* = 1*|x_V_, x_T_*) *> ξ*, the observer chooses *ŝ*_*C*=1_:

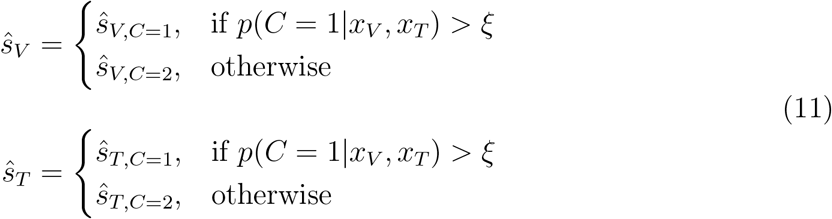

We evaluated the BCI framework against three alternative models, treating them as restricted or special cases of the full BCI model: (i) Forced Fusion, which assumes a common cause is always present (*p_common_* = 1); (ii) Full Segregation, which assumes signals are always independent (*p_common_* = 0); and (iii) Maximum Likelihood Estimation (MLE), a special case of forced fusion that assumes no informative spatial prior (effectively *p_common_* = 1 and *σ_P_ → ∞*). An overview of these models is presented in Table 1.

**Table 1:** Overview of models and their free parameters.

| Model | Free Parameters | # |
| --- | --- | --- |
| Bayesian Causal Inference (BCI) | $p_{common}, \mu_P, \sigma_P, \sigma_V, \sigma_T$ | 5 |
| Forced-fusion | $\mu_P, \sigma_P, \sigma_V, \sigma_T$ | 4 |
| Full-segregation | $\mu_P, \sigma_P, \sigma_V, \sigma_T$ | 4 |
| Maximum-likelihood estimation (MLE) | $\sigma_V, \sigma_T$ | 2 |

To account for the circular nature of the 360° environment while using a Gaussian BCI framework, we transformed all raw shot angles into a relative coordinate system. All visual stimulus locations (*s_V_*) were normalized to 0°, with tactile stimulus locations (*s_T_*) and participant responses (resp_*V*_, resp_*T*_) expressed as angular offsets ranging from *−*180° to +180°.

In this reference frame, the tactile stimulus *s_T_* was positioned at 0*^◦^, ±*45°, or *±*135° relative to the visual center.

Behavioral analyses were conducted across the full 360° space. BCI model fitting was restricted to the Front region because this region best isolated the intended manipulation of visual reliability: participants could use peripheral vision without first resolving the target through large exploratory head movements. By contrast, targets in the Side and Back regions often required overt orienting before the sensory judgment could be made. Accordingly, model fitting used only Front-region trials to estimate participant-specific parameters (*σ_V_, σ_T_, σ_P_, p_common_*), while regional effects were evaluated behaviorally.

We performed parameter estimation using the BCI Toolbox (Zhu et al., 2024). For each fit, we used the Bayesian Information Criterion (BIC) to identify the best-fitting decision strategy (Model Averaging, Selection, or Probability Matching) and to compare the full BCI model against the restricted alternative models (Forced Fusion, Full Segregation, and MLE). Because BIC favors parsimony, we also calculated McFadden’s pseudo-*R*^2^ 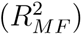 as an absolute goodness-of-fit measure based on the log-likelihood of the observed data (McFadden, 1974). For each subject and condition, we defined a null model by pooling data across all experimental conditions to generate a marginal probability distribution for each response channel. The null log-likelihood (*LL_null_*) was then compared to the log-likelihood of each candidate model (*LL_model_*). The final 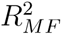 was defined as:

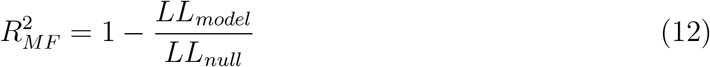

where a value of 0.2-0.4 indicates very good model fit.

## 3 Results

The results follow the three questions introduced above. First, does degraded vision increase tactile capture of visual responses? Second, do movement trajectories reveal action strategies that differ from the final perceptual endpoint? Third, does visual uncertainty change the tendency to bind visual and tactile cues, or does it primarily alter sensory noise and priors?

### 3.1 Accuracy

#### Unimodal Baselines

Participants were most accurate when the visual target was clearly visible (Accuracy = 0.96 *±* 0.08), least accurate when the visual target was hard to see (0.74 *±* 0.19), and showed intermediate performance when responding to vibration alone (0.90 *±* 0.07; Fig. 5 top row). A Linear Mixed Model (LMM) confirmed a significant main effect of visual reliability (*χ*^2^(2) = 373.12, *p* < .001). Performance was significantly better for high-reliability visual trials than for tactile-only trials (*z* = 4.65, *p <* 0.001), and tactile-only performance was significantly better than low-reliability visual performance (*z* = *−*13.91, *p <* 0.001).

**Figure 5:**
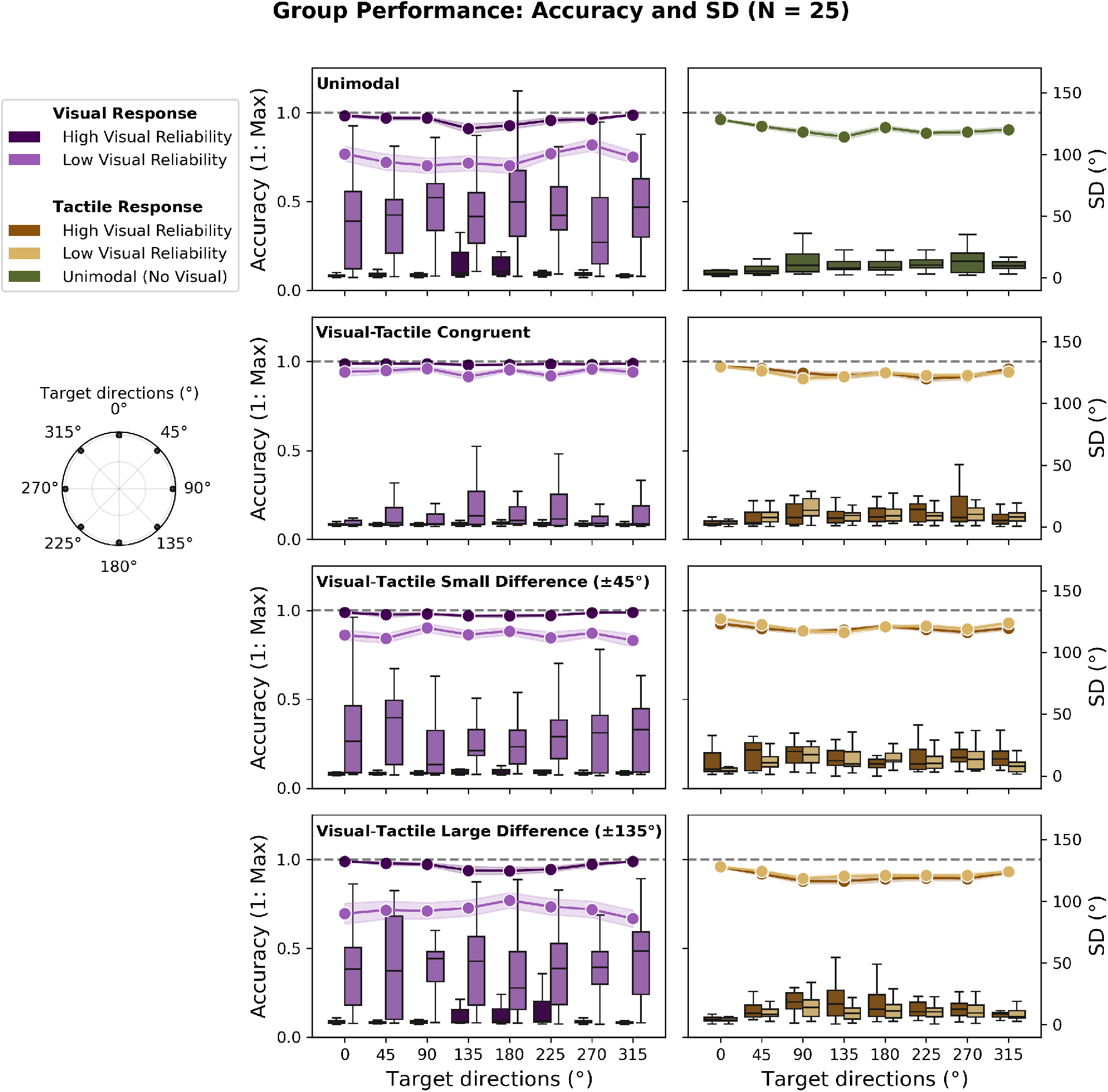
Group Performance (*N* = 25): Accuracy and Standard Deviation (SD) Across Visual Reliability Conditions. Line plots show mean accuracy ([0, 1]), and box plots show response standard deviation. Visual responses are shown in dark purple (high visual reliability) and light purple (low visual reliability). Tactile responses are shown in brown/yellow for high/low visual reliability, with green indicating unimodal tactile trials. The main pattern is that low visual reliability selectively reduces visual-response accuracy and increases variability.

#### Visual–tactile Integration under Spatial Disparity

When visual and tactile cues pointed to the same location, participants performed well regardless of which cue they were asked to report. Accuracy was high for both visual-response trials (High: 0.99 *±* 0.01; Low: 0.94 *±* 0.08) and tactile-response trials (High: 0.94 *±* 0.06; Low: 0.93 *±* 0.05).

The effect of visual reliability appeared when the visual and tactile cues pointed to different locations. When participants were asked to shoot the visual target, low visual reliability made performance worse as the conflict increased: accuracy dropped to 0.86 *±* 0.13 for small conflicts (VT-small) and 0.72 *±* 0.24 for large conflicts (VT-large). When the visual target was clearly visible, participants stayed accurate even during large conflicts (0.96 *±* 0.07).

Tactile-response trials showed a different pattern. When participants were asked to shoot toward the vibration, accuracy stayed high (0.90–0.92) even when visual reliability changed or the visual target conflicted with the tactile cue. A 3-way LMM confirmed this overall pattern, showing a significant Condition *×* Response Type *×* Reliability interaction (*χ*^2^(2) = 153.46, *p* < .001). The Response Type *×* Reliability interaction was also significant (*χ*^2^(1) = 8.98, *p* = .003), confirming that low visual reliability hurt visual responses more than tactile responses.

#### Effects of Spatial Region

We next asked whether performance changed depending on where the target appeared around the body: Front, Side, or Back (Fig. 3A). Overall, participants were most accurate for targets in front of them and slightly less accurate for targets to the side or behind them.

This pattern was already visible in the unimodal baseline trials. Tactile-only accuracy was higher for Front targets (0.93*±*0.06) than for Back targets (0.89*±*0.07; *z* = *−*2.20, *p* = .028). The LMM showed a significant main effect of Condition (*χ*^2^(2) = 171.13, *p* < .001) and a marginal effect of Region (*χ*^2^(2) = 5.70, *p* = 0.058). Importantly, Region did not interact with Condition (*χ*^2^(4) = 3.85, *p* = 0.43), meaning the same overall ordering held across space: high-reliability visual performance was best, tactile-only performance was intermediate, and low-reliability visual performance was worst.

The multisensory trials showed a similar pattern. Accuracy was highest for Front targets (*M* = 0.96 at the intercept) and declined slightly for Side (*z* = *−*0.040, *p* = .015) and Back targets (*z* = *−*0.044, *p* = .003). A 4-way LMM confirmed a significant effect of Region (*χ*^2^(2) = 10.45, *p* = .005) and a significant effect of Condition (*χ*^2^(2) = 14.36, *p* < .001). However, the full 4-way interaction between Condition, Response Type, Region, and Visual Reliability was not significant (*p* = .063). Thus, targets outside the front region were modestly harder to localize, but the main visual–tactile pattern was similar across the 360° space.

### 3.2 Directional Bias and Sensory Conflict

Directional bias was near zero when the visual target was clearly visible, and responses had low variance (*R >* 0.95; Fig. 6). When visual reliability was low, responses shifted toward the tactile cue, and this bias increased with visual–tactile conflict. Bias was 7.73° for small conflicts (*±*45°) and 11.24° for large conflicts (*±*135°). Large conflicts also produced more variable responses (*R* = 0.44).

**Figure 6:**
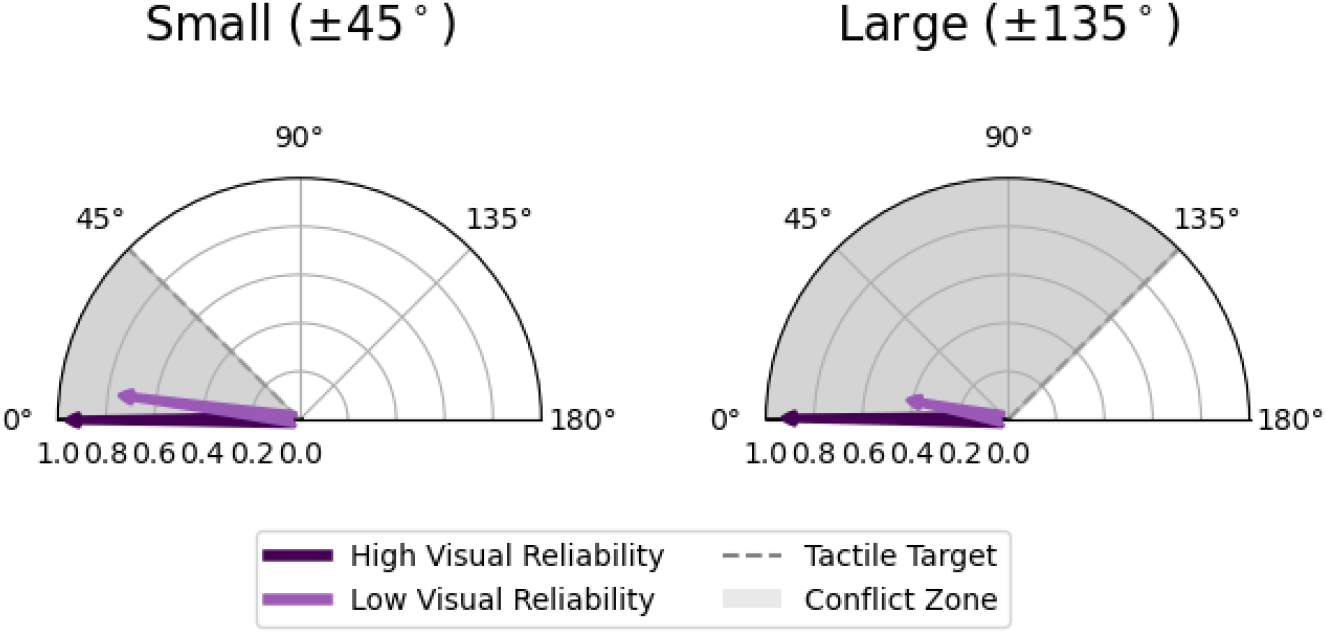
Mean Directional Bias and Response Precision across Spatial Disparity and Visual Reliability. Polar plots show aggregate response vectors for visual responses. The angular position of each arrow represents mean directional bias 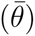, where 0° denotes the visual target and the gray shaded region represents the conflict zone toward the tactile target. Arrow length represents vector strength (*R*), a measure of circular precision ranging from 0 (random) to 1 (perfect consensus).

### 3.3 Movement Trajectories

Participants reached the target before shooting in most conditions, but they moved faster when responding to vibration than when searching for a visual target (Fig. 7). Tactile responses were fast and stable across conditions (mean RT *∼* 1.17 s). Visual responses were slower when the target was hard to see, with the low-reliability visual-only condition producing the longest reaction time (3.83 s). In multisensory trials, reaction times increased as visual–tactile conflict increased.

**Figure 7:**
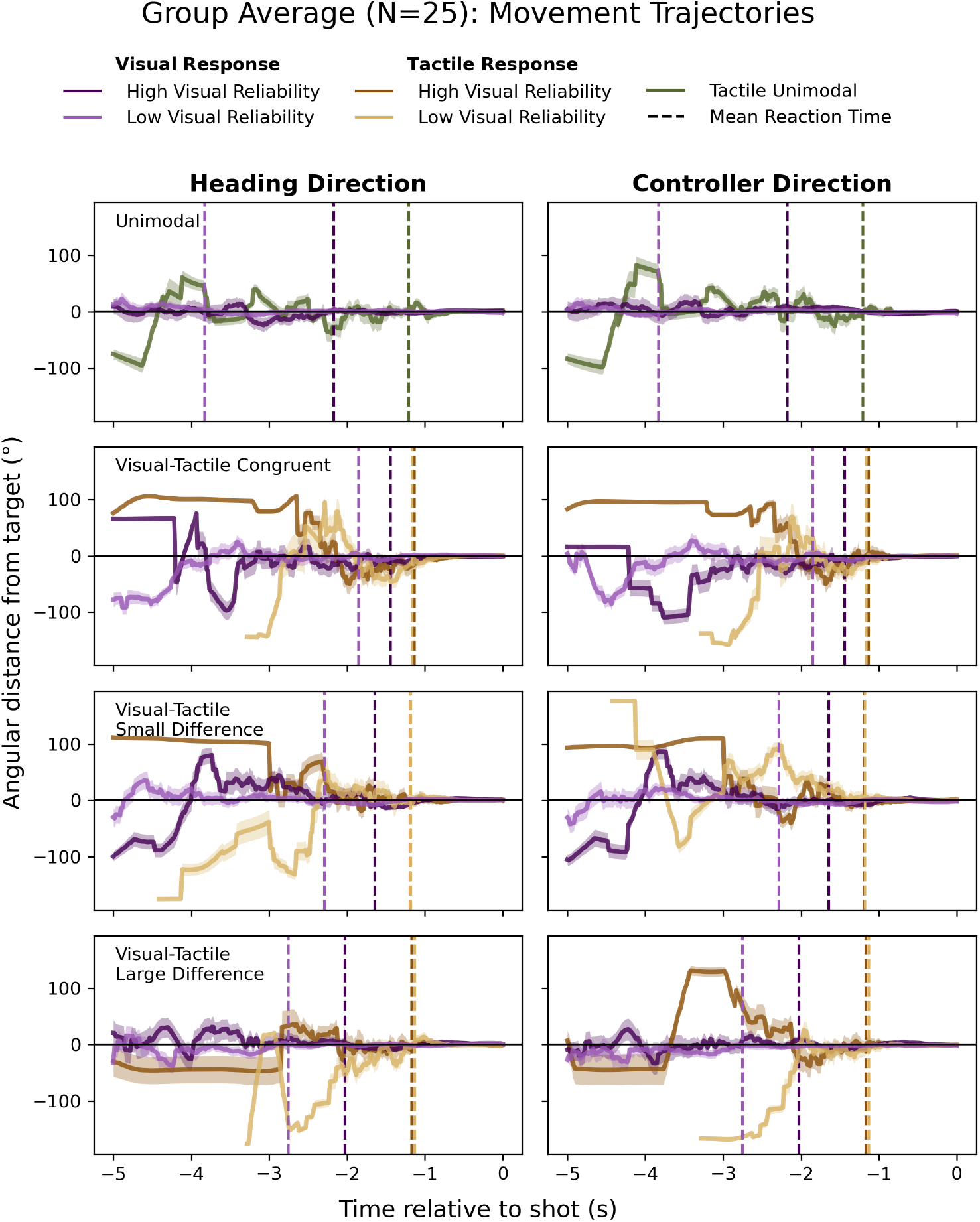
Group Average Movement Trajectories across Conditions and Visual Reliability Levels. Angular distance from the target (°) is plotted relative to shot time (0 s) for heading direction (left) and controller direction (right). Solid lines indicate group means (*N* = 25), shaded regions represent *±*1 SEM, and vertical dashed lines indicate mean reaction time for each condition and reliability level.

#### 3.3.1 Random Forest Classification

We used Random Forest classifiers to ask a simple question: could movement patterns reveal what kind of trial participants were doing? For each trial, we measured head and controller movement speed, acceleration, jerk, and path length. We then tested whether these movement features could classify task condition (Table 2; Fig. 8).

**Table 2:**
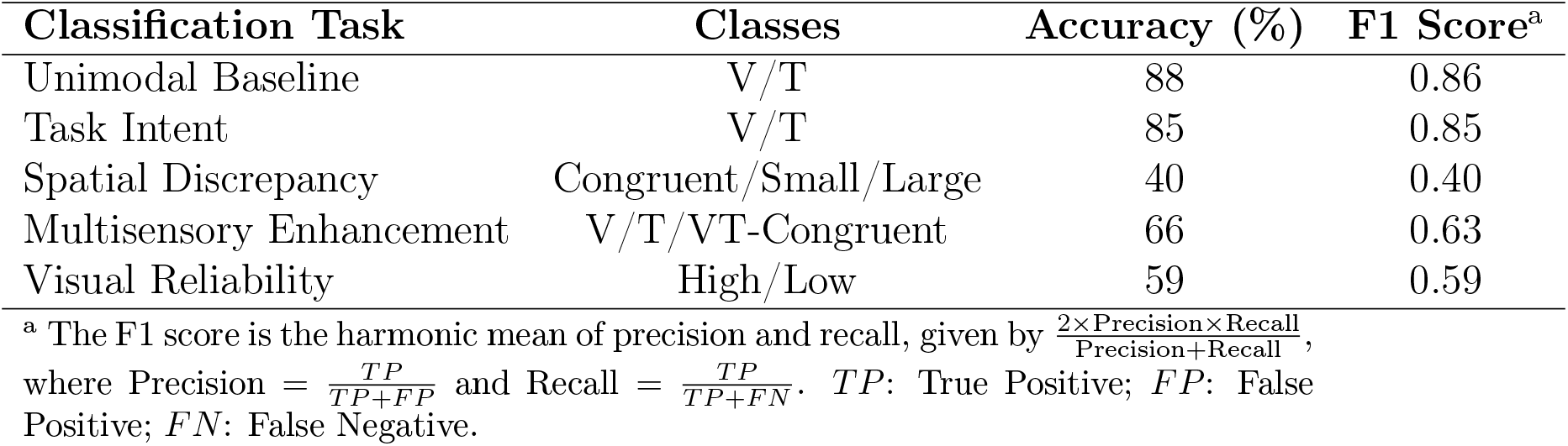
Performance Metrics for Random Forest Kinematic Classifiers. Classification accuracy and F1 scores are reported for each experimental dimension.

**Figure 8:**
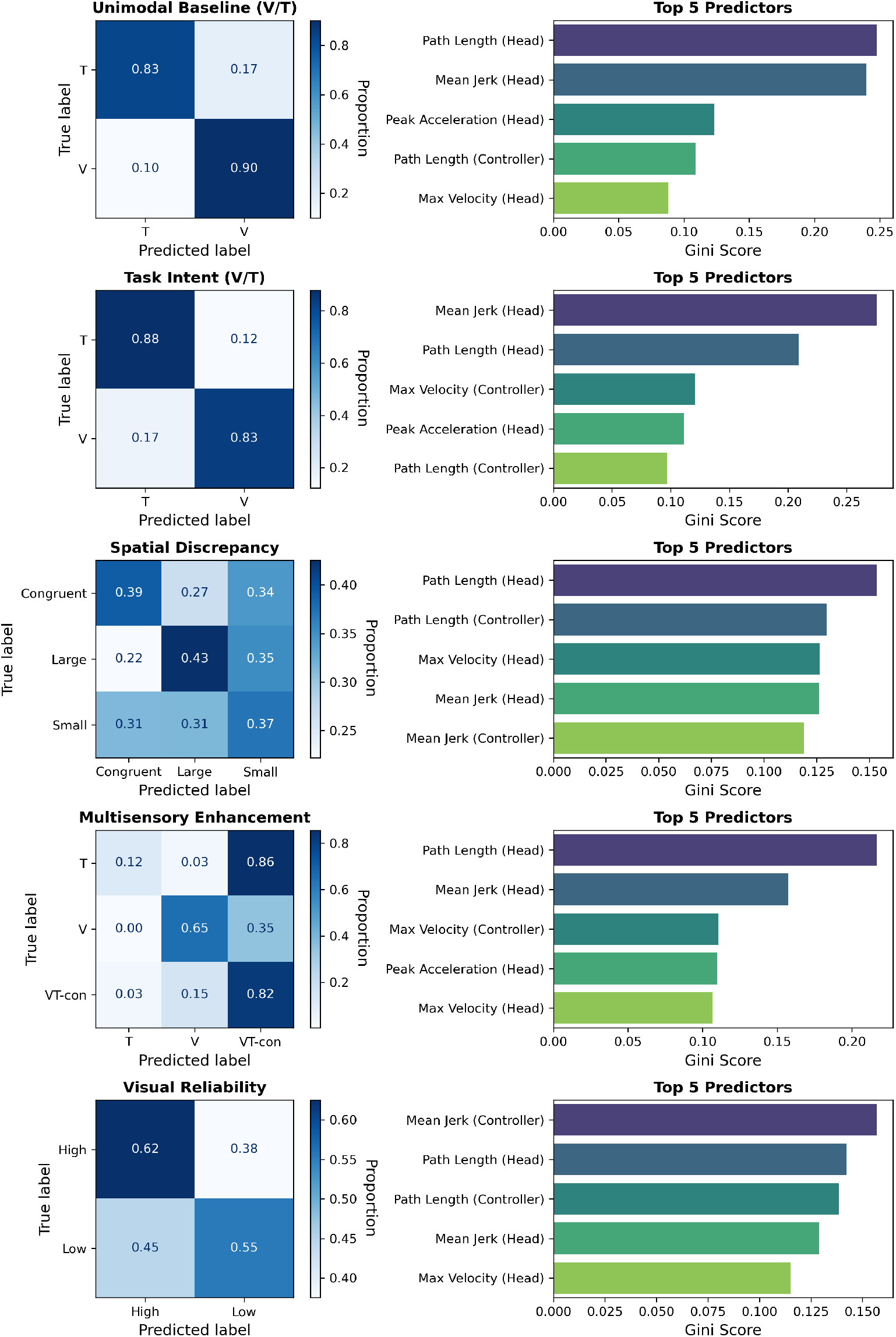
Random Forest Classification Performance and Kinematic Feature Importance. Each row represents a distinct classification task performed on movement kinematics (*N* = 25). **Left column:** normalized confusion matrices. **Right column:** top five predictors ranked by Gini Importance. Feature labels indicate whether data were recorded from the *Head* or *Controller* sensor.

The clearest result was that visual-guided and tactile-guided movements looked different. The classifier distinguished visual-only from tactile-only trials with 88% accuracy, and it identified whether participants were instructed to respond to the visual or tactile target in multisensory trials with 85% accuracy.

The classifier was much less accurate for the finer task details. It only weakly decoded the size of the visual–tactile conflict (40%; chance = 33.3%) and visual reliability (59%; chance = 50%). Thus, movement features clearly separated visual versus tactile guidance, but they were less sensitive to conflict size or visual reliability.

The multisensory enhancement analysis suggested that tactile guidance strongly shaped movement. Tactile-only trials were often classified as congruent visual–tactile trials (86% of the time; see Figure 8 fourth row), meaning that adding a matching visual target did not greatly change the movement pattern produced by vibration. Across classifiers, head path length and head jerk were among the most useful features, indicating that the efficiency and smoothness of head movement carried much of the task information.

### 3.4 Model Fitting

We next used computational models to ask what changed when vision became unreliable. The main model, Bayesian Causal Inference (BCI), assumes that the observer estimates whether visual and tactile cues came from the same source or from different sources. We compared BCI with three simpler models: Forced Fusion (FF), Full Segregation (FS), and Maximum Likelihood Estimation (MLE) (Ernst and Banks, 2002; Körding et al., 2007; Wozny et al., 2010).

We evaluated the models in two ways. Bayesian Information Criterion (BIC) asks which model explains the data with the fewest parameters; lower BIC is better. McFadden’s pseudo- *R*^2^ asks how well the model actually matches the observed responses; values typically range from 0 to 0.2 (good fit) up to 0.4+ (excellent fit).(McFadden, 1974).

#### 3.4.1 Model Comparison

When the visual target was clearly visible, BCI matched behavior best. It had the lowest mean BIC (141 *±* 4) and performed significantly better than FS (*p_adj_* = 0.015), MLE (*p_adj_ <* 0.001), and FF (*p_adj_ <* 0.001; Friedman Test: *χ*^2^ = 68.09, *p <* 0.001, post hoc pairwise comparisons with Bonferroni correction; Fig. 9 left). BCI also had the highest goodness-of-fit 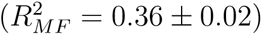.

**Figure 9:**
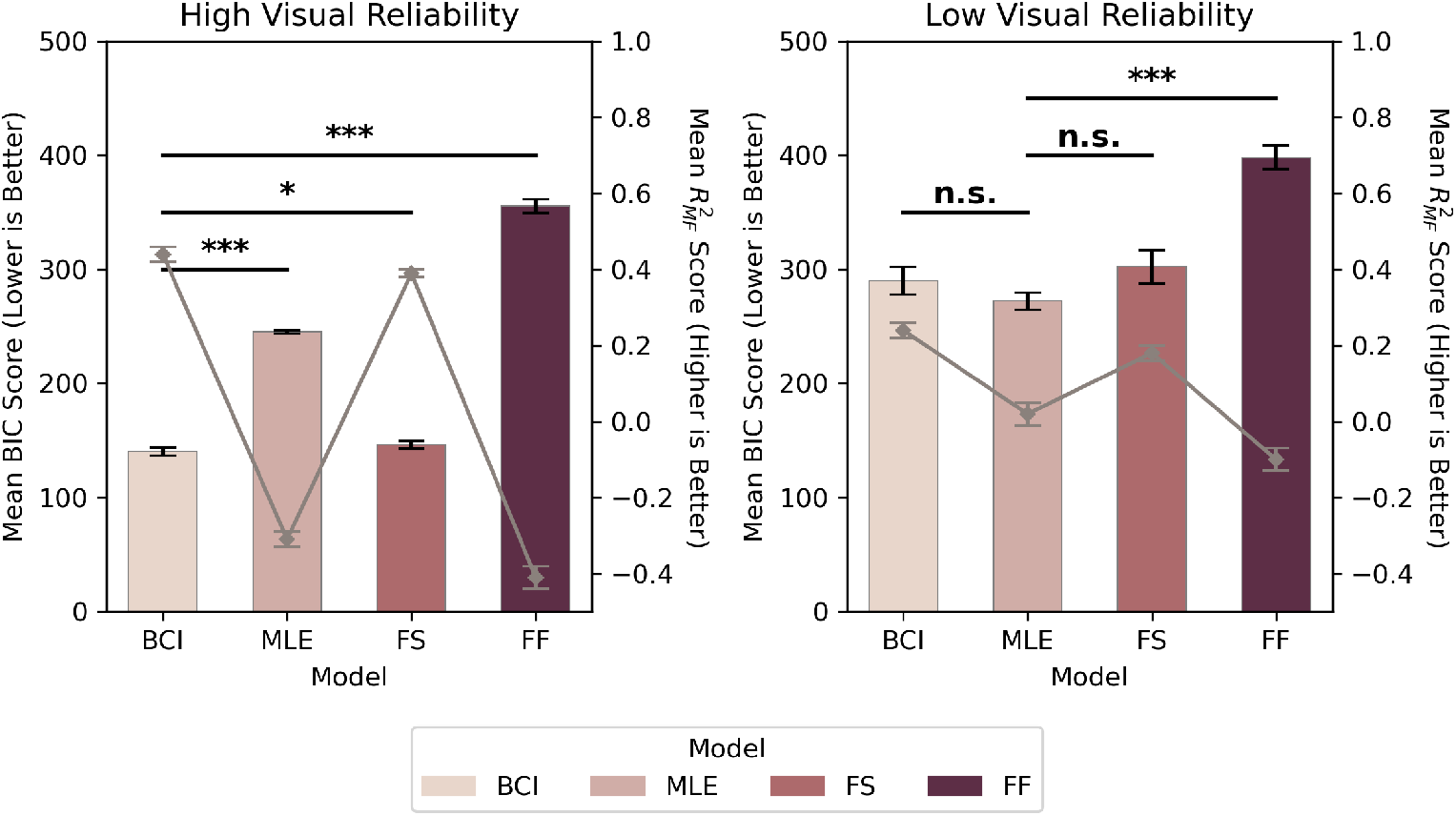
Model Comparison by Visual Reliability. Mean Bayesian Information Criterion (BIC) scores (bar plots, left axis; lower is better) and mean 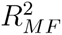 values (line plots, right axis; higher is better) are shown for BCI, MLE, FS, and FF models. Error bars represent *±*1 SEM. Asterisks denote statistical significance (n.s. = not significant, \**p <* 0.05, \*\**p <* 0.01, \*\*\**p <* 0.001).

When the visual target was hard to see, all models fit the data worse. MLE had a slightly lower mean BIC (272 *±* 8) than BCI (290 *±* 12), but this difference was not significant (*p_adj_* = 0.256). More importantly, MLE did not actually match the responses well: its 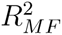 was near zero (0.02 *±* 0.02), while BCI still explained more of the data (0.22 *±* 0.01). In other words, MLE looked simpler, but BCI still described behavior better.

##### Model Fit and Accuracy

We also asked whether model fit was related to how well participants actually performed (Fig. 10). Under high visual reliability, this relationship was weak, likely because visual accuracy was already near ceiling. Under low visual reliability, however, participants with better accuracy also tended to have better BCI fits. BCI BIC scores were negatively correlated with accuracy for visual responses (*r* = *−*0.68, *p <* 0.001) and tactile responses (*r* = *−*0.63, *p <* 0.001). BCI 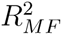 was also positively correlated with tactile accuracy (*r* = 0.52, *p* = 0.01), with a marginal trend for low-reliability visual accuracy (*r* = 0.39, *p* = 0.06). MLE did not show this relationship. Its BIC scores were not significantly related to observed accuracy (*r* = *−*0.09, *p* = 0.68), and its 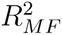 values were consistently lower than BCI. This suggests that MLE’s “better fit” under low visual reliability came mainly from being a simpler model, not from explaining behavior better.

**Figure 10:**
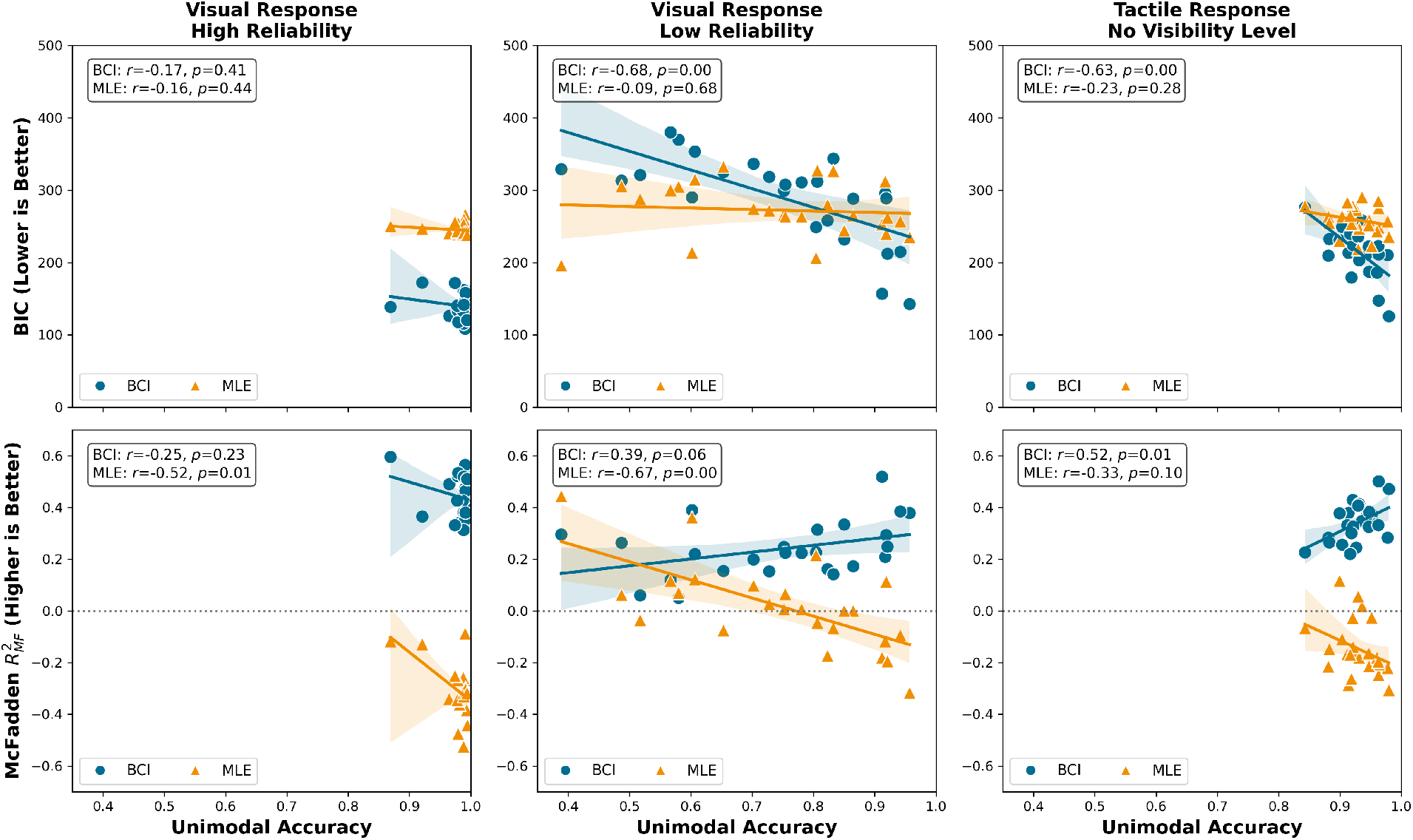
Relationship between Behavioral Accuracy and Model Fit. Each data point shows a participant’s mean visual or tactile accuracy plotted against BIC scores (lower is better) and 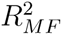 values (higher is better) from BCI and MLE model fits. Blue trend lines represent BCI; orange trend lines represent MLE.

#### 3.4.2 Parameter Estimation

Finally, we examined the fitted BCI parameters to ask what changed inside the model when vision became unreliable. The main change was visual uncertainty, not binding tendency (Table 3). Visual noise (*σ_V_*) increased strongly from high to low visual reliability (*t*(24) = *−*9.43, *p_adj_* < .001, *BF*_10_ = 7.5*e*6). The spatial prior also became broader under low reliability (*σ_P_* : *t*(24) = *−*3.55, *p_adj_* < .01, *BF*_10_ = 22.57), meaning the model estimated less certainty about where targets were expected to appear.

**Table 3:** Fitted BCI Parameters for Front Region. Mean parameter values *±* 1 standard error of the mean (SEM) are shown across high and low visual reliability conditions. Angular parameters (*σ_V_*, *σ_T_*, *σ_P_*, *µ_P_*) are expressed in degrees.

| Region | Parameter | High Reliability<br>(Mean ± SE) | Low Reliability<br>(Mean ± SE) | $t$ | $p_{adj}$ | $BF_{10}$ |
| --- | --- | --- | --- | --- | --- | --- |
| Front | $p_{common}$ | $0.42 \pm 0.05$ | $0.42 \pm 0.05$ | 0.04 | 1.00 | 0.21 |
| | $\sigma_P$ ($^\circ$) | $60.68 \pm 4.12$ | $84.49 \pm 5.03$ | -3.55 | <.01 | 22.57 |
| | $\mu_P$ ($^\circ$) | $48.77 \pm 1.18$ | $22.86 \pm 4.00$ | 6.48 | <.001 | 1.6e4 |
| | $\sigma_V$ ($^\circ$) | $3.14 \pm 0.38$ | $73.68 \pm 7.36$ | -9.43 | <.001 | 7.5e6 |
| | $\sigma_T$ ($^\circ$) | $27.50 \pm 2.54$ | $17.69 \pm 3.14$ | 2.35 | 0.14 | 2.07 |

Note: $p_{adj}$ reflects Bonferroni correction for five comparisons. $BF_{10} < 1$ indicates evidence favoring the null hypothesis.

The fitted prior mean (*µ_P_*) also differed between reliability levels, but this value should be interpreted cautiously. The prior distributions were very broad, so changes in the center of the prior had little influence on the final response. Group average BCI fits are shown in Fig. 11; corresponding MLE, FS, and FF fits are shown in Figs. A.3, A.4, and A.5.

**Figure 11:**
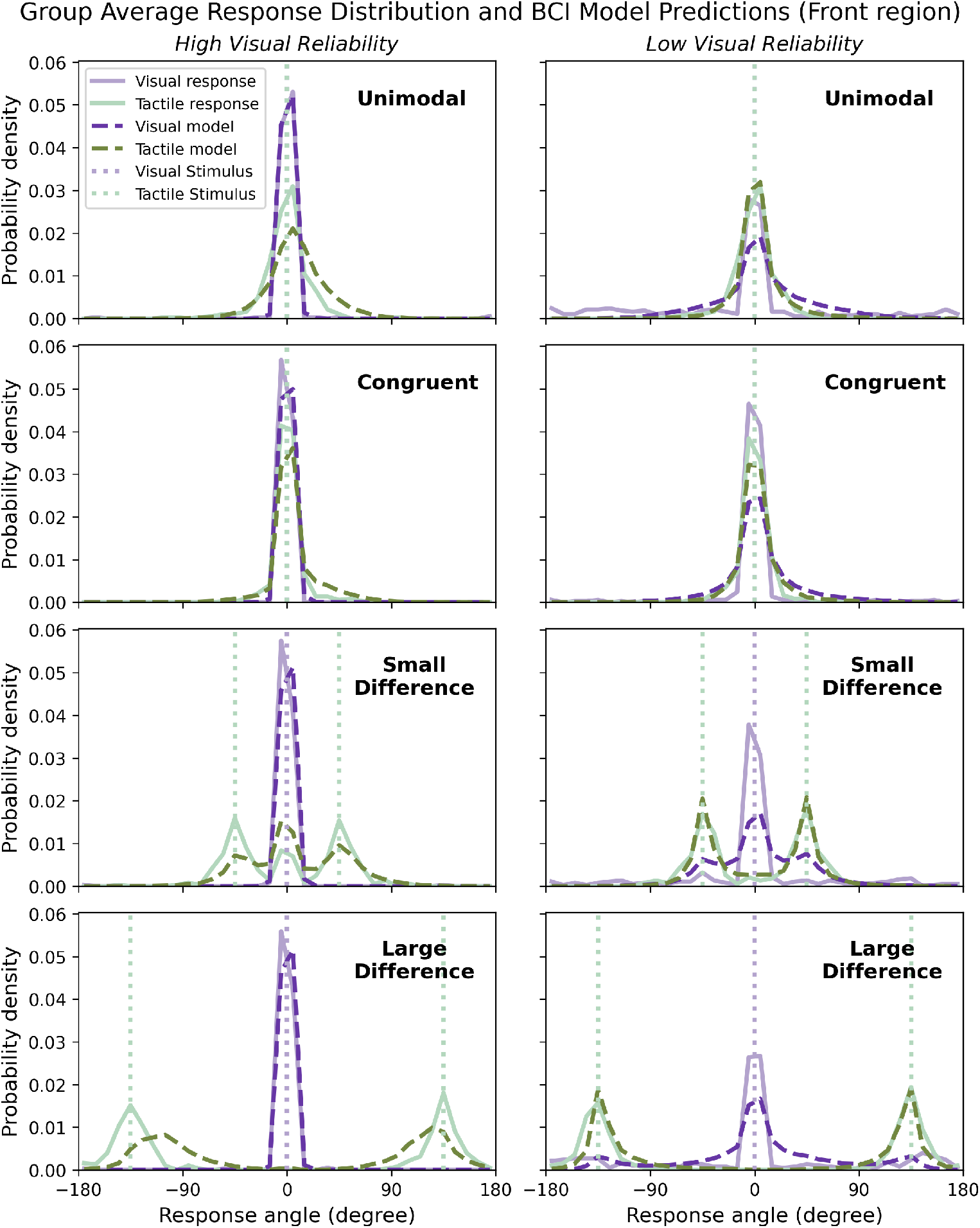
Group Average Bayesian Causal Inference Model Fits for the Front Region. Solid purple and green lines correspond to visual and tactile response data. Dashed purple and green lines correspond to model predictions for visual and tactile responses. Dotted lines indicate true stimulus values. All response data and stimulus locations are centered at 0°.

The tendency to bind visual and tactile cues did not change. The fitted *p_common_* was the same under high (0.42 *±* 0.05) and low (0.42 *±* 0.05) visual reliability (*t*(24) = 0.04, *p_adj_* = 1.00, *BF*_10_ = 0.21). Tactile noise (*σ_T_*) also did not differ significantly (*t*(24) = 2.35, *p_adj_* = 0.14, *BF*_10_ = 2.07). Thus, degraded vision mainly increased uncertainty in the visual signal; it did not make participants more likely to bind visual and tactile cues.

##### Model Identifiability and Parameter Recovery

We checked whether the model could recover its own parameters from simulated data (Appendix B, Figs. A.6, A.7). Recovery was strong for the main sensory parameters (*σ_V_* : *R*^2^ = 0.91; *σ_T_* : *R*^2^ = 0.63). Recovery was weaker for the prior parameters (*σ_P_*, *µ_P_*), which is expected because the fitted priors were broad and had limited influence on responses. Overall, these checks support the main parameter result: visual reliability changed sensory uncertainty, while the binding tendency remained stable.

## 4 Discussion

This study asked three questions about visual–tactile integration in an active 360° VR task. First, does degraded vision increase tactile capture of visual responses? Second, do movement trajectories reveal action strategies that differ from the final response? Third, does degraded vision change the tendency to bind visual and tactile cues, or does it mainly change sensory uncertainty? The results answer these questions in a consistent way: degraded vision made visual responses more vulnerable to tactile cues, tactile signals strongly shaped movement, and the tendency to bind cues remained stable while visual uncertainty increased.

### Degraded Vision Increased Tactile Capture

When the visual target was clearly visible, participants showed little bias toward the tactile cue and their responses were precise. When the visual target was hard to see, responses shifted toward the tactile cue, and this shift increased as the visual–tactile conflict became larger. Large conflicts also produced more variable responses. This shows that tactile cues had the strongest influence when vision was unreliable.

This pattern was specific to visual responses. When participants were asked to respond to vibration, performance stayed relatively stable even when a visual target was also present. In other words, unreliable vision made visual localization more vulnerable to touch, but conflicting visual information did not strongly disrupt tactile localization. Across the full 360° space, targets outside the front region were modestly harder to localize, but the main pattern (stronger tactile influence when visual reliability is low) was similar across Front, Side, and Back regions.

### Tactile Cues Strongly Shaped Movement

Movement trajectories showed that tactile cues influenced action in a way that was not fully captured by final accuracy alone. Participants moved faster when responding to vibration than when searching for a visual target. In multisensory trials, reaction times increased as visual–tactile conflict increased, suggesting that conflict slowed the process of selecting or confirming a response.

The Random Forest analysis supported the same conclusion. Movement features clearly distinguished visual-guided from tactile-guided trials, but they were less sensitive to conflict size or visual reliability. Tactile-only trials were often classified as congruent visual–tactile trials, suggesting that adding a matching visual target did not greatly change the movement pattern already produced by vibration. This points to a simple division of labor: tactile cues may help guide the early orienting movement, while vision helps refine the final target location (Paillard, 1996; Woodworth, 1899; Tagliabue and McIntyre, 2014).

### Binding Tendency Stayed Stable

The modeling results suggest that degraded vision changed uncertainty, not the basic tendency to bind visual and tactile cues. When the visual target was clearly visible, BCI matched behavior best. When the visual target was hard to see, all models fit worse, but BCI still described behavior better than the simpler MLE model. MLE sometimes looked favorable by BIC because it had fewer parameters, but it did not match the observed responses well and did not track accuracy across participants.

The fitted BCI parameters clarify why. Low visual reliability strongly increased visual noise (*σ_V_*), and the spatial prior became broader, meaning the model estimated greater uncertainty about where targets were likely to appear. In contrast, *p_common_* stayed the same across high and low visual reliability. Thus, participants did not compensate for poor vision by becoming more likely to bind visual and tactile cues. Instead, the binding tendency remained stable, while visual uncertainty increased.

### Implications and Limitations

Together, these findings suggest that visual–tactile integration in active VR depends strongly on sensory reliability. Degraded vision does not appear to change the basic tendency to bind cues, but it makes visual responses noisier and more vulnerable to tactile influence. At the same time, tactile signals provide a fast body-centered cue for action. This makes tactile displays useful even when they are relatively low-resolution, because they can support rapid orienting before vision refines the final response (Lloyd-Esenkaya et al., 2020; Caraiman et al., 2019; Spence, 2014).

Several limitations remain. BCI model fitting was restricted to the Front region, so the present modeling results do not show whether the same parameters would explain Side and Back targets. Future work should test circular BCI models that can fit the full 360° space directly. Models with lapse parameters may also better capture occasional search failures under very low visibility. Finally, this study used static targets; future work should test whether the same tactile-guided movement pattern holds during dynamic navigation.

## 5 Conclusion

This study shows that visual–tactile integration in 360° VR depends strongly on how reliable vision is. When the visual target was clearly visible, participants localized it accurately with little influence from touch. When the visual target was hard to see, responses shifted toward the tactile cue, especially when the visual and tactile cues were far apart.

Movement data showed a complementary pattern. Tactile cues supported fast, stable orienting movements, whereas visual responses slowed down when the target was degraded or conflicted with touch. This suggests that tactile signals can guide the body quickly, while vision helps refine the final target location.

The BCI modeling showed that degraded vision mainly changed uncertainty, not binding tendency. Visual noise increased and spatial expectations became broader, but *p_common_* stayed stable across high and low visual reliability. In other words, participants did not become more likely to bind visual and tactile cues when vision was poor; instead, they relied less on visual information.

Together, these findings suggest that tactile cueing can support action in visually uncertain environments without requiring highly precise spatial information. A wearable tactile display may be especially useful for navigation because it can provide fast body-centered guidance when visual information is limited, noisy, or slow to find.

## Data and code availability statement

The figures in this article, as well as the data and plotting scripts necessary to reproduce them, are available openly under the CC-BY license (Chan, 2026).

## Acknowledgements

A.C. thanks the Croucher Foundation for a scholarship.

## A Supplemental Information

**Figure A.1:**
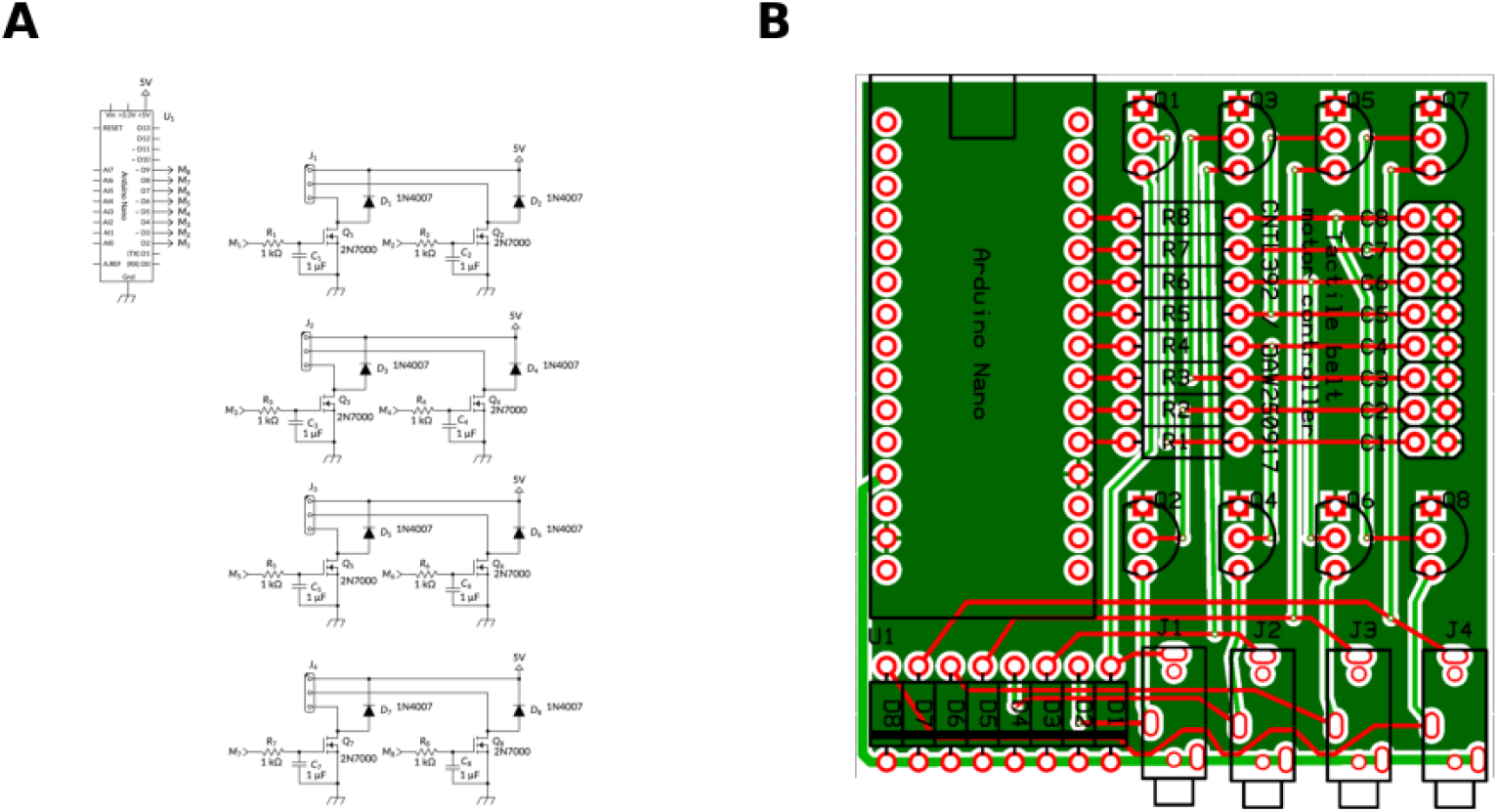
Detailed circuit schematics and PCB layout.

**Figure A.2:**
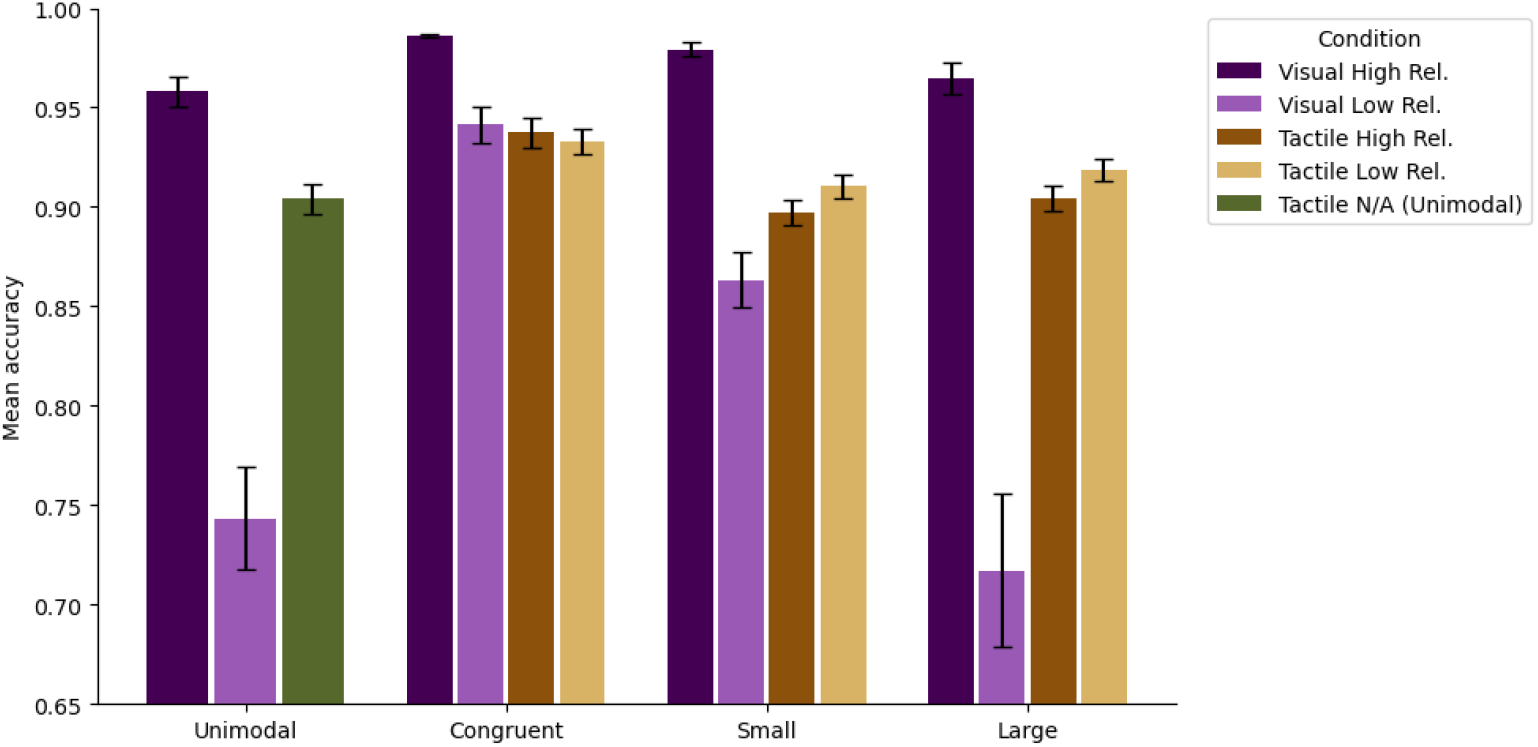
Mean accuracy across conditions, all directions pooled. Error bars represent *±*1SEM.

**Figure A.3:**
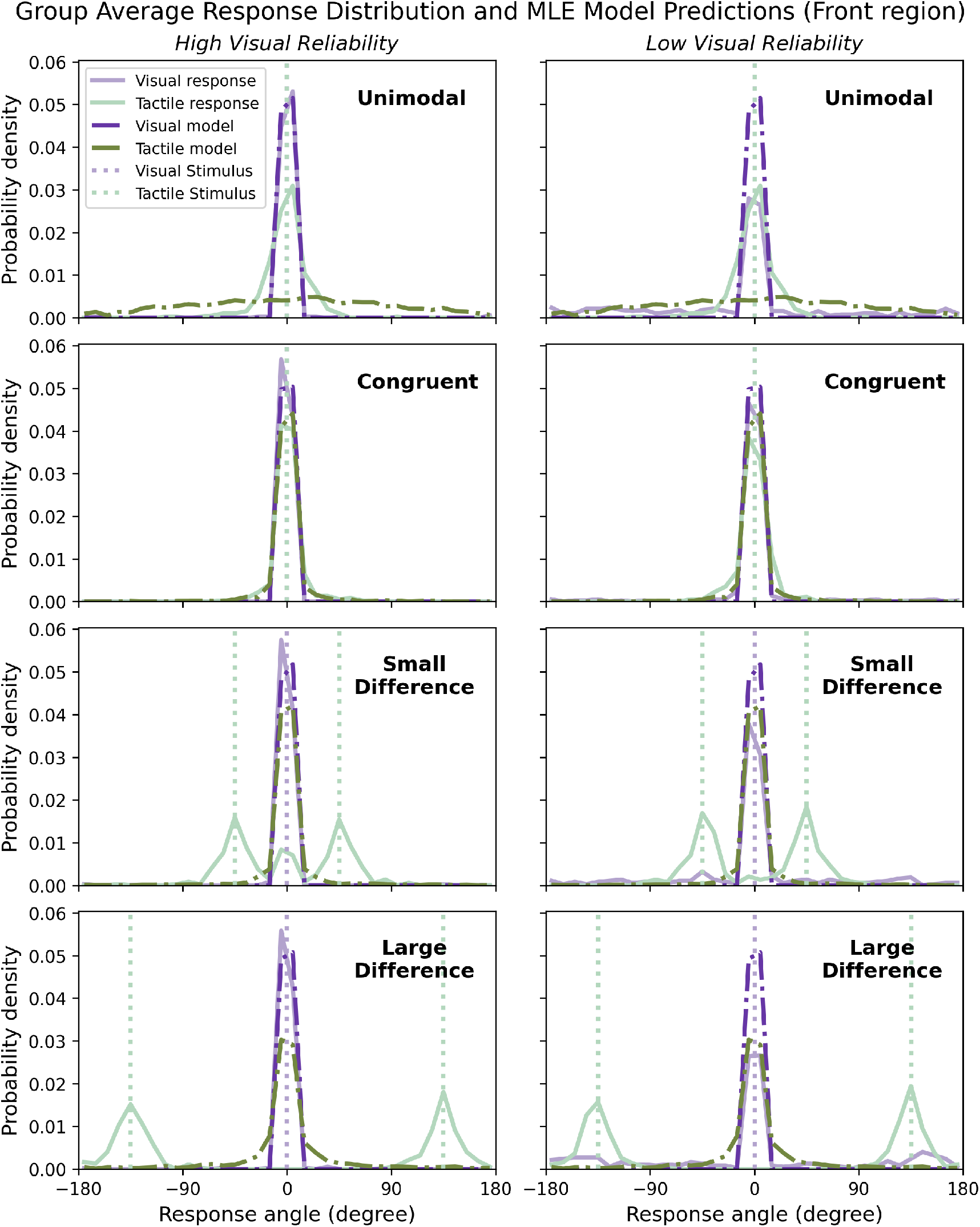
Group average MLE model fits. Solid purple and green lines correspond to the visual and tactile response data, dashed purple and green lines correspond to model predictions for visual and tactile responses, and dotted purple and green lines correspond to true stimulus values. All observed response data and stimulus locations are centered at 0°.

**Figure A.4:**
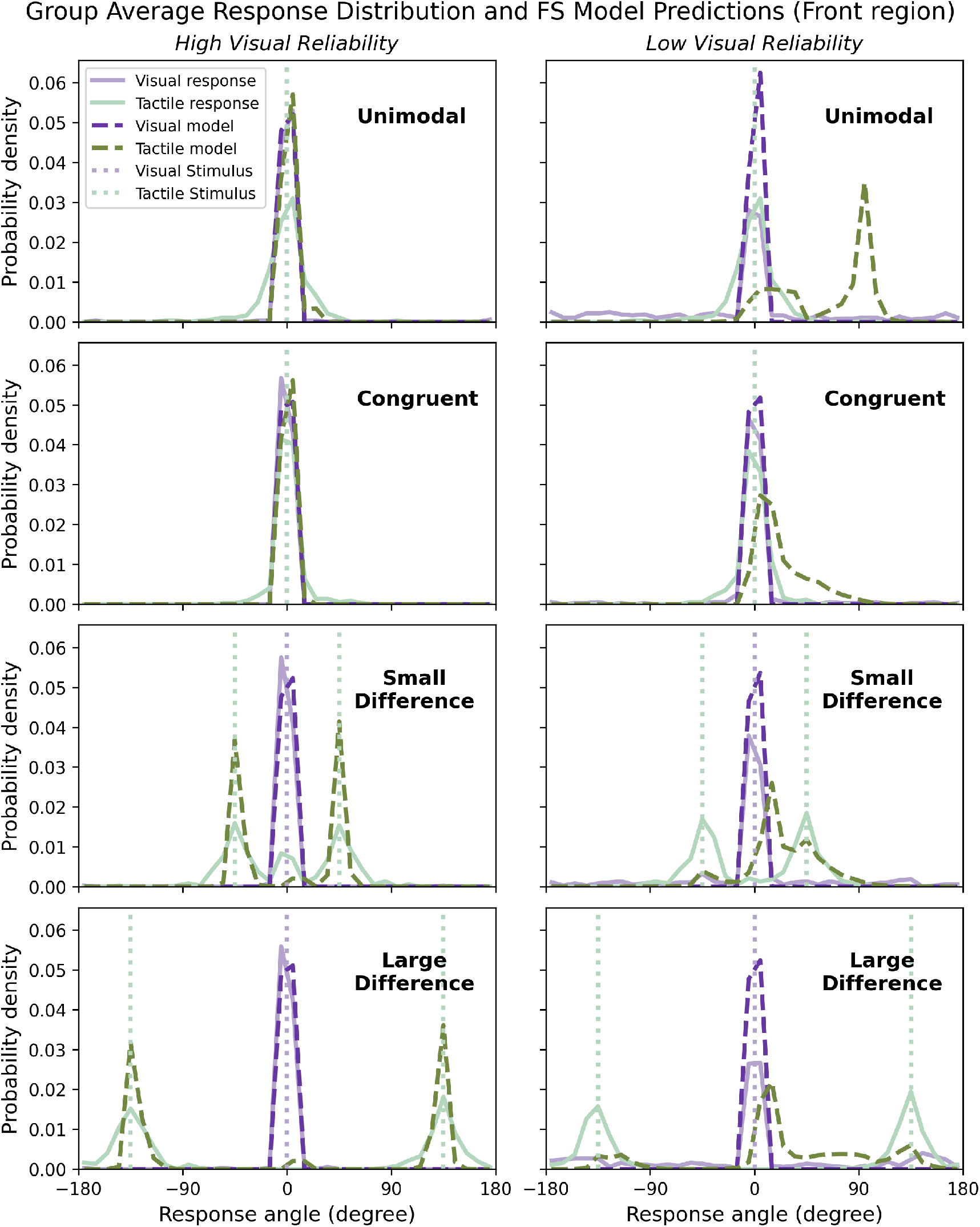
Group average FS model fits. Solid purple and green lines correspond to the visual and tactile response data, dashed purple and green lines correspond to model predictions for visual and tactile responses, and dotted purple and green lines correspond to true stimulus values. All observed response data and stimulus locations are centered at 0°.

**Figure A.5:**
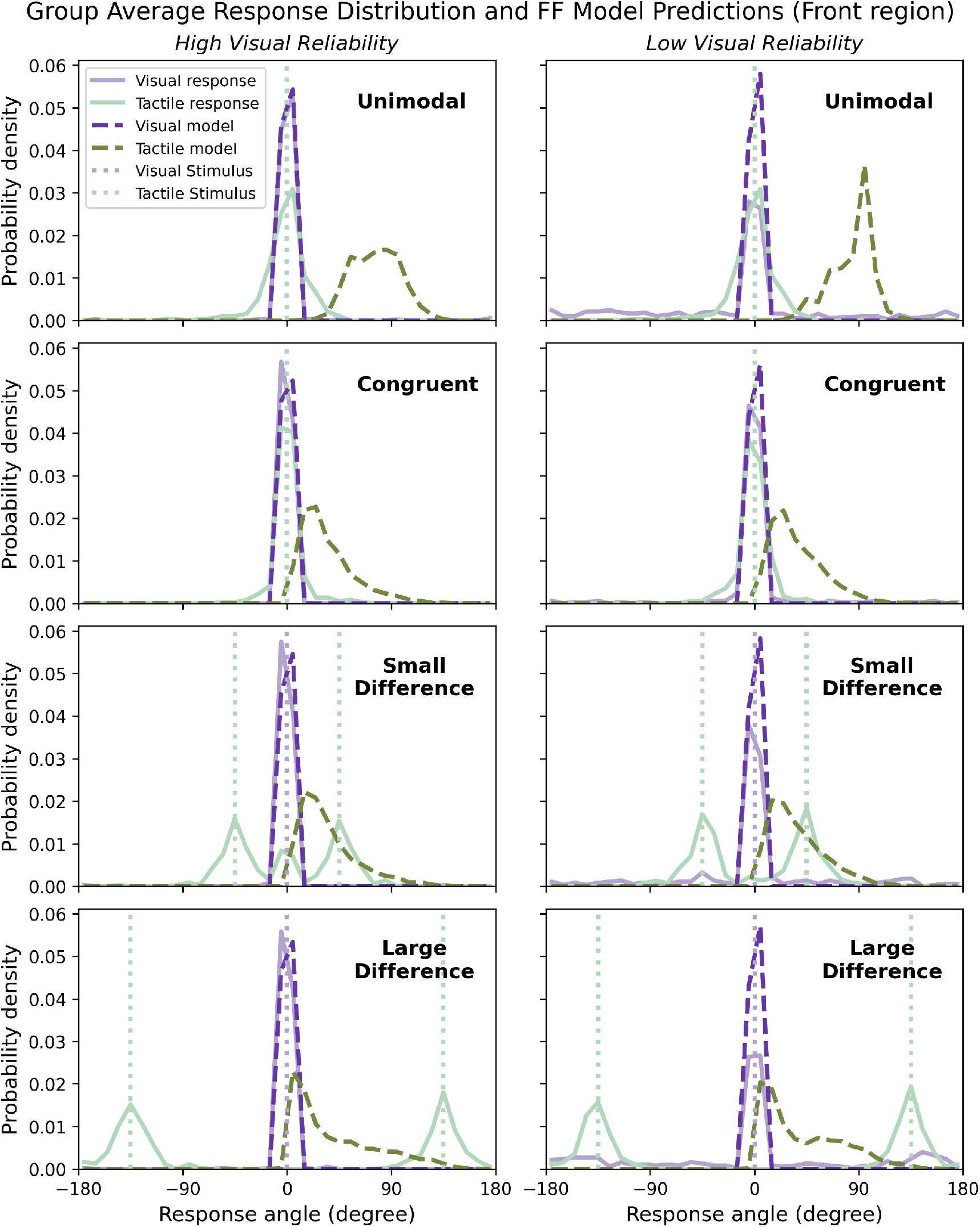
Group average FF model fits. Solid purple and green lines correspond to the visual and tactile response data, dashed purple and green lines correspond to model predictions for visual and tactile responses, and dotted purple and green lines correspond to true stimulus values. All observed response data and stimulus locations are centered at 0°.

**Figure A.6:**
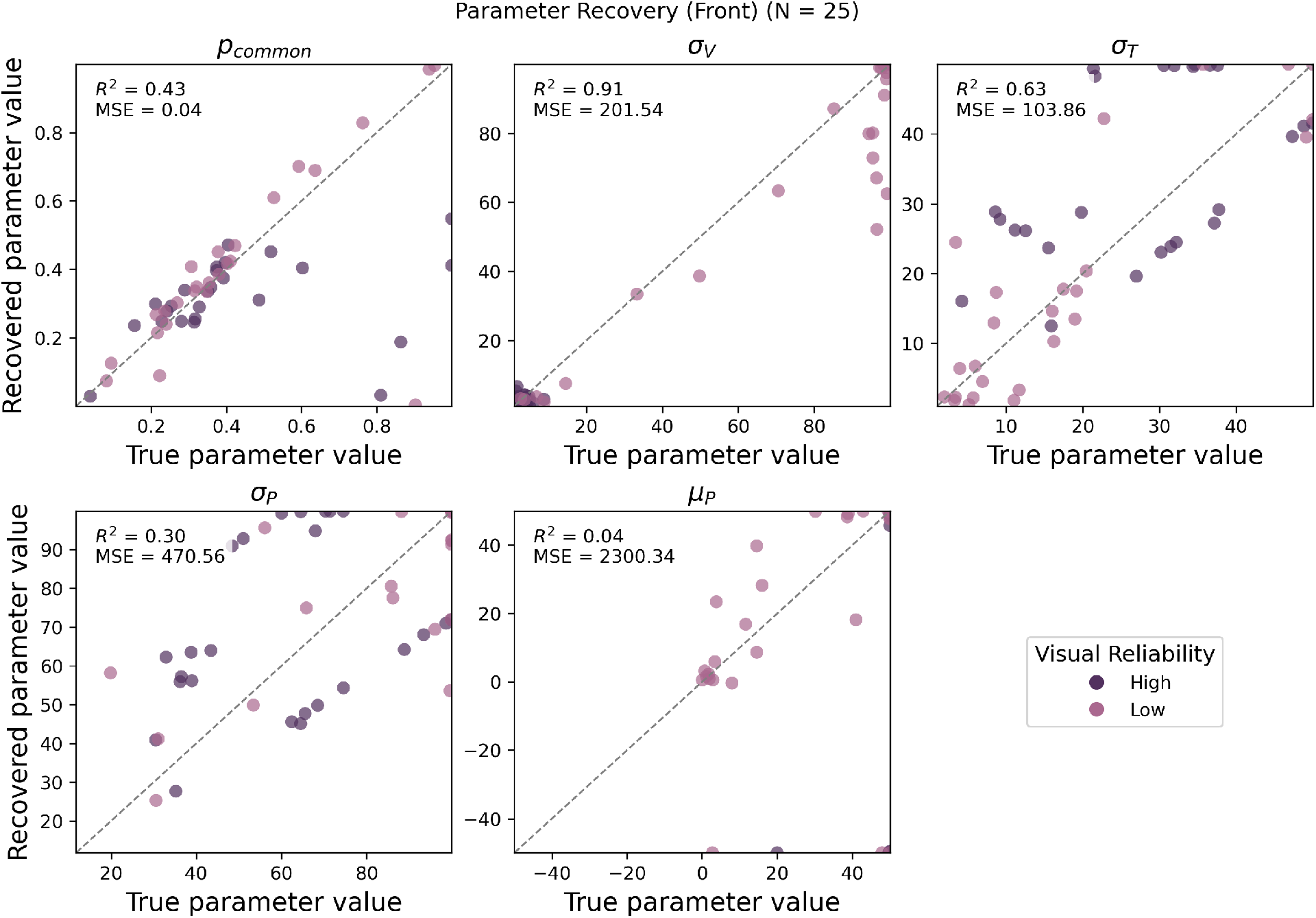
Parameter recovery analysis for the Bayesian Causal Inference (BCI) model. Scatter plots illustrate the correspondence between ground-truth and recovered parameter values across 25 synthetic datasets. To evaluate model identifiability and experimental robustness, synthetic data were generated using each participant’s estimated parameters and subsequently re-fitted. In each panel, recovered values are plotted against ground truth, with dashed gray lines representing the identity (*y* = *x*). Sensory likelihood parameters (*σ_V_* and *σ_T_*) show strong recovery, with *σ_V_* achieving an *R*^2^ = 0.91. While *p_common_*displays a lower *R*^2^ (0.43), its low Mean Squared Error (MSE = 0.04) indicates that estimates are accurate but clustered within a narrow range, making *R*^2^ sensitive to minor deviations. The weaker recovery of spatial prior parameters (*σ_P_, µ_P_*)–characterized by low *R*^2^ and high MSE–is expected, as the fitted priors were sufficiently broad to be weakly informative.

**Figure A.7:**
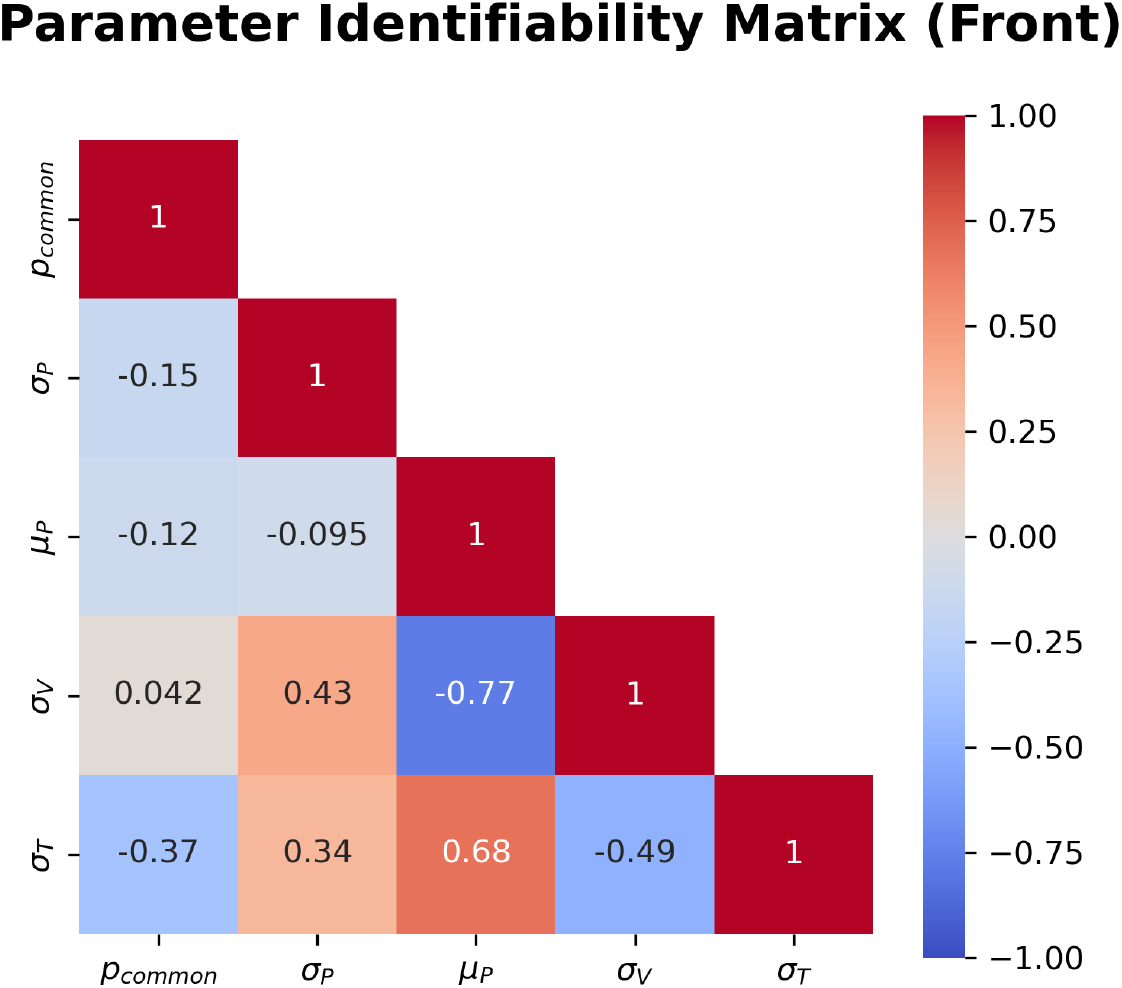
Parameter Identifiability Matrices. Pearson correlation coefficients (*r*) calculated for the Front region (*N* = 25). The primary parameters (*p_common_*, *σ_P_*, *σ_V_*) exhibit minimal inter-parameter correlation, confirming that the BCI framework uniquely identifies these values regardless of spatial location. Observed correlations between visual noise (*σ_V_*) or tactile noise (*σ_T_*) and the spatial prior mean (*µ_P_*) are considered functionally negligible. Given the uninformatively broad spatial prior (*σ_P_*) and the limited recoverability of *µ_P_*, these spatial offsets do not meaningfully influence the resulting perceptual estimates, nor do they compromise the stability of the core causal and sensory parameters.

